# Neuromuscular Basis of *Drosophila* Larval Rolling Escape Behavior

**DOI:** 10.1101/2023.02.01.526733

**Authors:** Patricia C. Cooney, Yuhan Huang, Wenze Li, Dulanjana M. Perera, Richard Hormigo, Tanya Tabachnik, Isuru S. Godage, Elizabeth M.C. Hillman, Wesley B. Grueber, Aref A. Zarin

## Abstract

When threatened by dangerous or harmful stimuli, animals engage in diverse forms of rapid escape behaviors. In *Drosophila* larvae, one type of escape response involves C-shaped bending and lateral rolling followed by rapid forward crawling. The sensory circuitry that promotes larval escape has been extensively characterized; however, the motor programs underlying rolling are unknown. Here, we characterize the neuromuscular basis of rolling escape behavior. We used high-speed, volumetric, Swept Confocally-Aligned Planar Excitation (SCAPE) microscopy to image muscle activity during larval rolling. Unlike sequential peristaltic muscle contractions that progress from segment to segment during forward and backward crawling, the muscle activity progresses circumferentially during bending and rolling escape behavior. We propose that progression of muscular contraction around the larva’s circumference results in a transient misalignment between weight and the ground support forces, which generates a torque that induces stabilizing body rotation. Therefore, successive cycles of slight misalignment followed by reactive aligning rotation lead to continuous rolling motion. Supporting our biomechanical model, we found that disrupting the activity of muscle groups undergoing circumferential contraction progression lead to rolling defects. We use EM connectome data to identify premotor to motor connectivity patterns that could drive rolling behavior, and perform neural silencing approaches to demonstrate the crucial role of a group of glutamatergic premotor neurons in rolling. Our data reveal body-wide muscle activity patterns and putative premotor circuit organization for execution of the rolling escape response.

**Significance Statement:** To escape from dangerous stimuli, animals execute escape behaviors that are fundamentally different from normal locomotion. The rolling escape behavior of Drosophila larvae consists of C-shaped bending and rolling. However, the muscle contraction patterns that lead to rolling are poorly understood. We find that following the initial body bending, muscles contract in a circumferential wave around the larva as they enter the bend, maintaining unidirectional rolling that resembles cylinder rolling on a surface. We study the structure of motor circuits for rolling, inhibit different motor neurons to determine which muscles are essential for rolling, and propose circuit and biomechanical models for roll generation. Our findings provide insights into how motor circuits produce diverse motor behaviors.

## Introduction

Early in the evolution of animals, nervous systems specialized to permit locomotion [1]. While locomotion supports multiple aspects of evolutionary success (*e.g.* allocating resources, finding mates), one of the most critical of these is escape: the transformation of sensory input into motor output to avoid imminent danger [2–4]. Escape behaviors are rapid and stereotyped yet must be flexible enough to allow animals to evade multiple sources of harm and readjust when danger subsides[3, 5, 6]. Escape behaviors across species often differ fundamentally from exploratory locomotion[3, 7–11]. This specificity suggests that dedicated neural circuits or unique activity patterns within shared locomotor circuits are employed during escape. While many studies have investigated how sensory input promotes escape [2, 3, 6, 8, 9, 12–16], the neuromuscular activity that generates escape movements have been characterized in few model organisms[8, 12, 17, 18]. Furthermore, the model systems in which escape movement generation has been studied have yielded limited understanding of how the sensory circuits that promote escape drive motor circuits. By characterizing escape motor circuits in the *Drosophila* model, with its well-studied sensory system and nearly complete connectome, we aim to understand how sensory input is transformed into motor output during escape.

The *Drosophila* larval body consists of twelve segments, with abdominal segments containing up to 60 different muscles [19]. Forward crawling consists of sequential segmental contractions that propagate from posterior to anterior segments and engages all muscles [19–21]. Upon experiencing harmful mechanical stimuli or heat, larvae initiate a nocifensive escape consisting of C-shaped bending, rolling, and rapid forward crawling [22]. Rolling causes fast lateral motion—which is faster than escape crawling alone, and can dislodge attacking parasitoid wasps [23, 24]. This behavior is initiated by activity of class IV (cIV) dendritic arborization neurons, polymodal nociceptors that tile the body wall [23, 25]. Several downstream partners of cIV neurons have been identified and reconstructed using serial transmission electron microscopy [26–30]. Activation of any of several interneurons that are downstream of cIV neurons is sufficient to evoke a rolling escape response [26, 27, 30], but how these interneurons drive downstream motor activity patterns remains poorly characterized.

Despite progress in understanding nociceptive circuitry, characterizing neural and muscular activity during escape behavior presents challenges. In contrast to crawling, rolling behavior is asymmetric, with larvae rolling laterally in one direction. However, both the larval body and central nervous system (CNS) are symmetric on either side of the dorsal and ventral midlines. The hemisegment unit is important to consider during rolling behavior, since the bilaterally symmetric neural and muscle activity that occurs during crawling must be broken during rolling, setting up a fundamental difference between these two behaviors.

In this study, we examine the muscle activity and motor circuits responsible for escape bending and rolling using a combination of high-speed 3D imaging of fluorescent calcium indicators expressed in muscles, functional manipulations of motor circuits, and connectomics approaches. We compare our findings to the motor activity that drives crawling to identify what features of peristaltic locomotor drive are preserved in escape, and what motor features are unique to rolling escape locomotion. Both behaviors involve sequential motor activity and antagonistic drive of distinct muscle groups, but our results highlight fundamental differences in the motor patterns. In particular, muscle contractions progress circumferentially around the larva during rolling, in contrast to the anteroposterior progression of muscle contractions during crawling. These data provide a foundational view of motor activity during larval rolling, narrowing in on a full sensory to motor understanding of an escape behavior.

## Results

### Muscle activity patterns in rolling escape behavior

*Drosophila* larval rolling escape is comprised of C-shaped bending followed by lateral rolling [22]. Rolling can be triggered experimentally by activation of nociceptive sensory neurons or by central neurons including the Goro command neuron [26]. We confirmed that in response to optogenetic activation of Goro or a global heat nociceptive stimulus, larvae engage in bending and rolling behavior. We found that larvae can bend to the left or right, and, independent of bend direction, may roll in a clockwise or counterclockwise direction (**Figure 1A; Movie S1**). Thus, upon optogenetic activation of nociceptive circuitry, larvae can engage four distinct, yet related, escape motor patterns.

**Figure 1:**
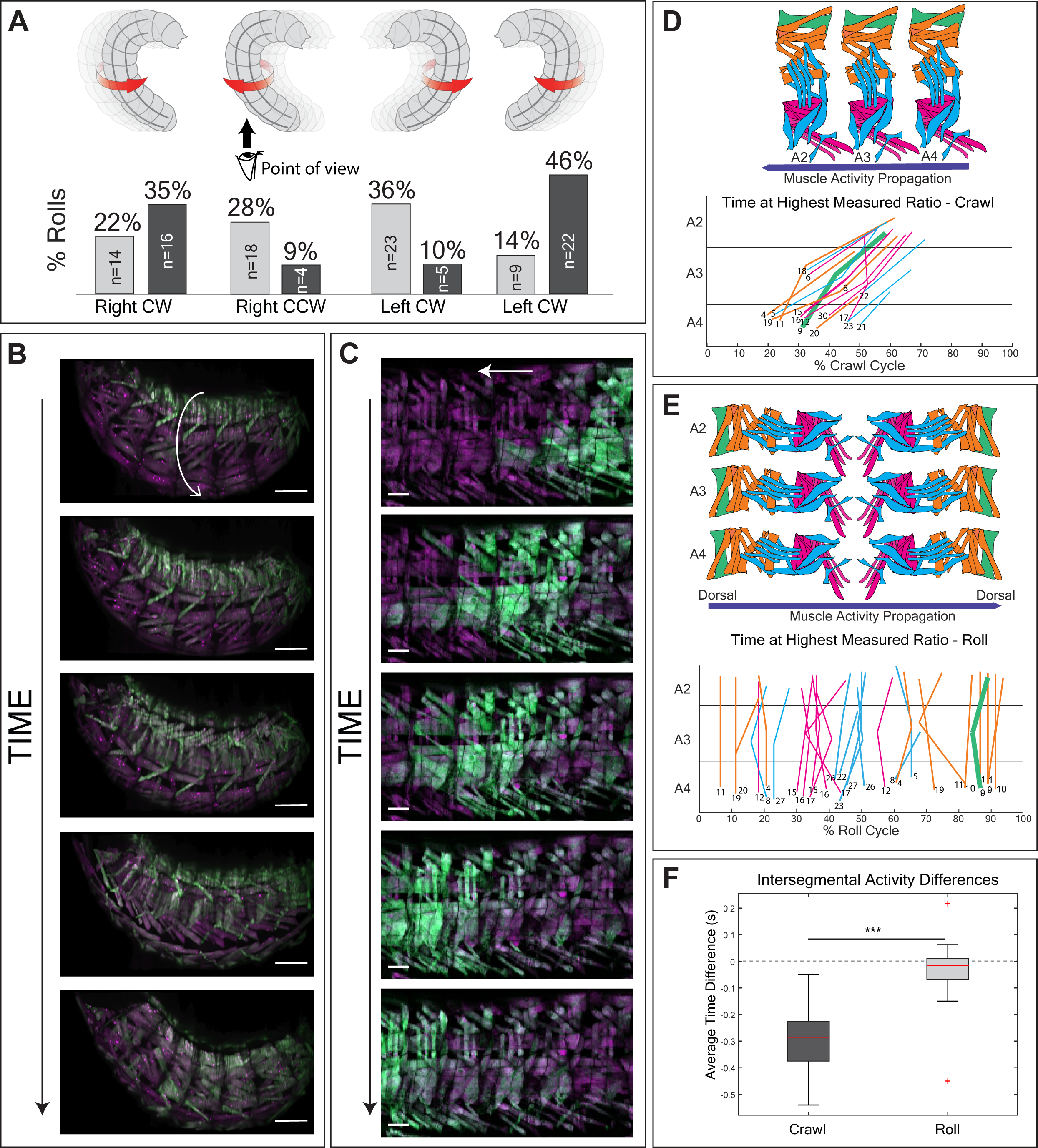
Muscles demonstrate a segmentally synchronous, circumferential wave of activity during escape. (**A**) Schematic illustrating four patterns of escape observed, based on combination of which side larva bends toward (left or right) and which direction the larva rotates (clockwise or counterclockwise). Translation direction is determined by direction of rotation. Histogram shows frequency of escape patterns observed in response to sustained optogenetic Goro activation (light gray, n = 64 rolls) or in response to global heat + vibration stimulus (dark gray, n = 47 rolls). (**B**) SCAPE dual-color stills from single roll bout, showing same muscle appearing in focal plane simultaneously across segments. Muscle GCaMP increased primarily on the bent side of the larva. Arrow indicates direction of larval rotation. (**C**) Confocal dual-color stills from single crawl bout in larva, demonstrating increase in GCaMP brightness from posterior to anterior muscles during single crawl bout. (**D**) Schematic of muscle arrangement in three neighboring hemisegments, color coded by muscle groups, as established in Landgraf *et al.*, 2003. Blue arrow indicates posterior-to-anterior propagation of muscle activity. Peak ratiometric muscle fluorescence times during single crawl bout across three segments with muscle number at bottom of line. During forward crawling, muscles of segment A4 reach peak activity before A3, and muscles of segment A3 reach peak activity before A2 segment, demonstrating the propagation of peristaltic contraction from posterior to anterior segments. Representative homologous muscle across hemisegments and peak activity lines in green for clarity of segmental propagation of activity during crawling. (**E**) Three-segment schematic of hemisegments, color coded by muscle groups, as established in Landgraf *et al.*, 2003. Blue arrow indicates circumferential propagation of muscle activity. Highest observed ratiometric muscle fluorescence times during single roll cycle with SCAPE across segments A2-A4 with muscle number at bottom of each line. Same muscle types across segment A2 to A4 simultaneously reach their peak activity. Muscles are color-coded according to panel **D**, demonstrating dorsal to ventral to dorsal (circumferential) propagation of muscle contraction. Representative homologous muscle across segments and peak activity lines in green indicate an example of segmental synchrony of activity during rolling. (**F**) Comparison between time difference of muscles in segments A2-A4 for forward crawling versus rolling (crawl: n = 2 crawls, 2 larvae, 86 muscles; roll: n = 4 rolls, 3 larvae, 372 muscles). Negative values indicate that muscles in the adjacent posterior segment are active before muscles in the adjacent anterior and “0” indicates synchronous contraction. Mann-Whitney U tests were performed between intersegmental roll lag values and intersegmental crawl lag values. P values are indicated as ***p<0.001. Scale bars = 100µm (B), 50µm (C).

Circuitry triggering the rolling escape behavior is well-studied in *Drosophila* larvae, but how circuitry converges on premotor and motor systems is not known. We therefore sought to determine the muscle activity that underlies the escape rolling motor pattern. We imaged larvae using Swept Confocally-Aligned Planar Excitation (SCAPE) microscopy, a volumetric imaging technique that permits high-speed, high-resolution, 3D imaging of behaving animals [31–38]. We induced rolling using Goro activation in larvae expressing mCherry and GCaMP6f in all body wall muscles. We resolved activity of individual muscles along the entire length of the larva and approximately half of the body thickness, at 10 volumes per second (**Figure 1B**; **Figure S1A,B; Movie S2)**. We predicted that as GCaMP6f/mCherry ratios increased, muscle length would decrease, reflecting muscle contraction upon activation. Indeed, these two measurements showed an inverse relationship, suggesting that GCaMP6f/mCherry ratios can be used as an indicator of muscle contraction (**Figure S1C**). We focused the bulk of our analysis on muscles in mid-segments A2-A4 since activity in these showed the greatest dynamic range (**Figure S1C**). We analyzed roughly 19 muscles per hemisegment in A2-A4 across multiple roll events, and in A1 and A5 during two roll events, totaling over 520 muscle measurements.

SCAPE movies revealed that muscles are most active along the bent side of the larva, consistent with a role for asymmetric muscle contractions in C-shaped bending (**Movie S3**; **Figure 1B**). Ratiometric calcium signals for many muscles tended to decrease as muscles moved out of the bend (**Movie S3; Figure S2A-D, Figure S3A-C**) and to increase as muscles rotated into the bend (**Movie S3; Figure S2E**). To contrast escape rolling and crawling at the level of individual muscles, we compared SCAPE imaging data collected during rolling to previously acquired confocal data on muscle activity during crawling [21] (**Figure 1B,C, Figure S3D,E**). As expected, measurements of muscle peak activity during crawling revealed a delay between muscle contraction in neighboring segments during peristalsis (**Figure 1C,D**). By contrast, during rolling, segmentally homologous muscles showed synchronous contractions, primarily on the side of the larva entering the bend. Also, in contrast to peristalsis, sequential muscle activity traveled around the circumference of the larva during rolling (**Figure 1 B,E,F, Figure S4, Figure S3D,E**). Notably, we found that while dorsal (D) and ventral (V) longitudinal and oblique muscles demonstrated significantly greater activity along the bent side than the stretched side of the larva, lateral transverse (LTs) and ventral acute (VAs) muscles show a different activation pattern. Specifically, VA muscles show only a moderate decrease in activity between the bent and stretched sides of the larva. LT muscles show heterogeneous activity patterns, where ∼60% demonstrate roughly equivalent activity in each phase of the roll and ∼40% demonstrate increasing activity as they approach the stretched side of the larva (**Figure 2A-C**). These primary features of muscle activity are consistent across segments A1-A5, though LT muscles are more likely to increase activity as they approach the stretched side of the larva in midsegments A2-A4 than in distal segments A1 and A5, and VA muscles show significant decreases in activity in A1 and A5, analogous to longitudinal muscles (**Figure S3B-D**). Therefore, LT and VA muscles do not follow the typical circumferential wave of activity seen in other body wall muscles

**Figure 2:**
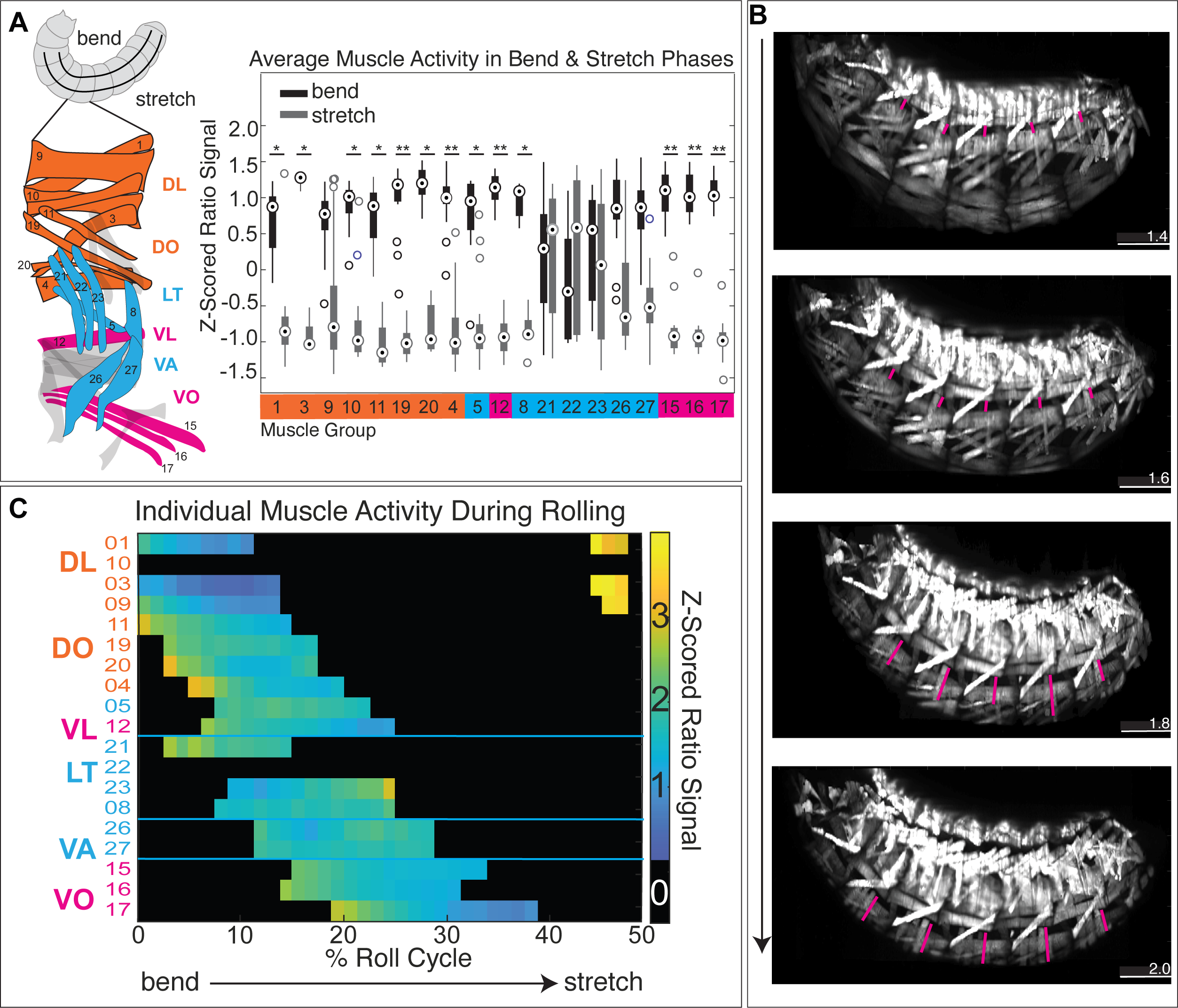
Intrasegmental muscle activity patterns demonstrate functional antagonism during escape. (**A**) Hemisegment schematic of example measured muscles color-coded according to Landgraf *et al.*, 2003 (*left*). Boxplot showing mean z-scored ratio signal for individual muscles from frames when muscles were along the bent side of the larva (black) vs. along the stretched side of the larva (gray) (*right*). Data are grouped and color-coded along x-axis according to dorsoventral order and similarity of activity patterns. Orange muscles and magenta muscles (dorsal longitudinal, DL; dorsal oblique, DO; ventral longitudinal, VL; ventral oblique, VO) show increased ratio signal along the bent side of the larva, while cyan (lateral transverse, LT) muscles show on average equivalent ratiometric signal on bent and stretched sides. Cyan (ventral acute, VA) muscles show elevated activity on bent side, but the difference between activity in bent and stretched sides is insignificant. (n = 3 rolls, 3 larvae, 280 muscles). See **Figure S2F** for frequency of equivalent vs. increasing LT and VA activity patterns. (**B**) Example ratiometric SCAPE stills of larval escape. Magenta lines highlight lateral transverse (LT) muscles, demonstrating low ratio signal while rotating out of the bend and increased ratio signal while rotating toward the stretched side of the larva. (**C**) Heatmap of z-scored ratio signal across individually measured muscles in one example hemisegment, organized from dorsal (top row) to ventral (bottom row). LT ratiometric traces show mixed activity patterns, (23 increases activity while 21 and 8 show roughly equivalent activity as they rotate toward the bend), an activity pattern different from other muscles. VA ratiometric traces show roughly equivalent activity throughout the roll. Color bar to the right of shows range of z-scored ratio signal values. Black indicates frames when muscles were out of the FOV and not measured. Scale bars = 100µm.

Altogether, these data demonstrate crucial distinctions between motor patterns during rolling and crawling: 1) muscle activity during rolling is synchronous across segments but is intersegmentally asynchronous during crawling; 2) muscle activity is left-right asymmetric during rolling but is left-right symmetric during crawling; 3) rolling involves progression of muscle contractions around the circumference of the larva, while crawling involves progression of muscle contractions along the anteroposterior axis. As an important exception to (2) above, we predict that as larvae roll, there are short periods of bilateral synchronicity (momentary left-right hemisegmental symmetry), during which homologous muscles flanking the dorsal (i.e., left and right DLs) or ventral midline (i.e., left and right VOs) enter the bend and co-contract. However, rolling and crawling are similar in that the within-segment muscle activity patterns both demonstrate opposing functions of longitudinally-spanning versus transverse-spanning (LT and VA) muscles.

### The biomechanics of larval rolling motion

So far, we have shown that activation of Goro command neuron results in propagation of muscle contraction around the circumference of the larvae. But how does the circumferential progression of muscle contraction lead to the rolling motion that translates larval body on the surface? To address this question and further elucidate the biomechanics of larval rolling, we propose the following mechanics model for inward (counterclockwise with respect to the head) (**Figure 3A-E** and **Movie S4)** and outward (clockwise with respect to the head) (**Figure 3F-I** and **Movie S4**) rolling motions.

**Figure 3.**
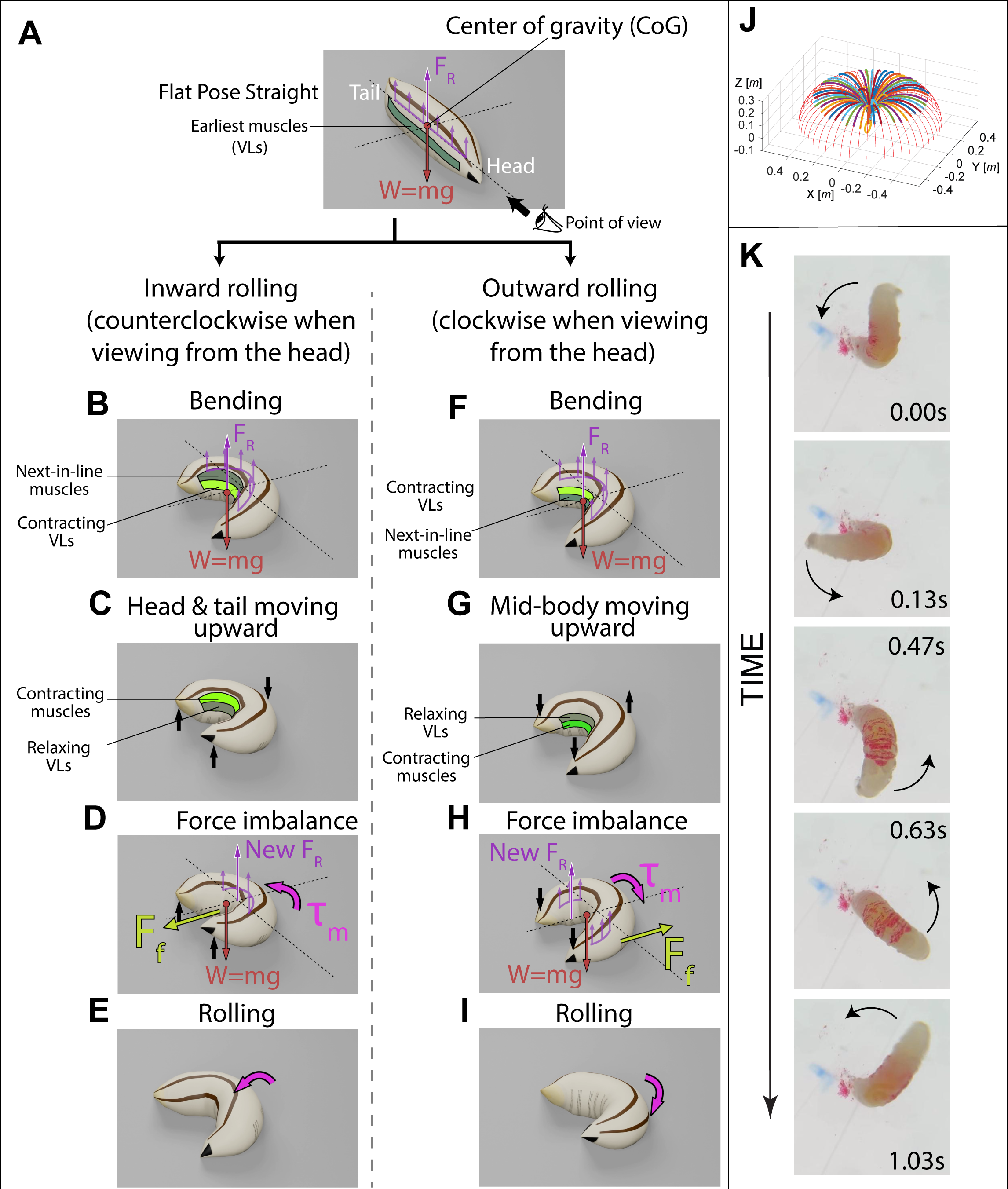
3D model of larva demonstrating the biomechanics of rolling. Larval rolling is driven by the discrete and infinitesimal propagation of the bending plane and the subsequent force and moment imbalance. There are two types of rolling: inward rolling (**B-E)** is when the larva rolls toward the bend (counterclockwise when viewing from the head), and outward rolling (**F-I)** is when the larva rolls away from the bend (clockwise when viewing from the head). **A)** Model of a larva lying flat on a surface without bending. The weight (*W=mg*) is balanced by and aligned with the sum of ground reaction forces (*F_R_*) that are evenly distributed at contact points between the larval body and the surface. *m* is the mass of the object and *g* gravitational acceleration. **B) and F)** Bending is the initial step of both types of rolling. The larva bends into a C-shape by contracting its ventrolateral muscles (VLs). This causes a shift of its center of gravity (CoG) toward the bend and away from the anteroposterior axis of the body, but W is still aligned with the sum of ground reaction force (F_R_). **B-E)** Steps specific to inward rolling (counterclockwise with respect to the head). **B)** After initial bending, the muscles above VLs are the next in line to be activated. **C)** The muscles above the VLs are activated, causing the head and tail to slightly move upward, leading to a slight shift of the bending plane. **D)** Because the head and tail are no longer touching the ground, the point of application of the sum of ground reaction forces moves to the middle body segments that are in contact with the ground. The concentrated sum of reaction forces to the mid-segments no longer aligns with its body weight. The imbalance of forces causes a moment (torque), τ_m_, which induces counterclockwise rolling. While this torque makes the larva tend to slip away from the bend, a static friction (F_f_) is applied to the larva, oriented toward the bend to prevent slipping. **E)** The larva rotates counterclockwise (towards the bend opening) as it falls back to the surface. **F-I)** Steps specific to outward rolling (clockwise with respect to the head). **F)** After initial bending, the muscles below VLs are the next to be activated. **G)** The muscles below VLs are activated, causing a downward movement of the head and tail while the mid-segments of the body are slightly lifted. **H)** The change of contact points causes the sum of F_R_ to move away from the middle segments toward the bend, no longer aligning with weight. The static friction Ff in this case is oriented away from the bend. **I)** The imbalance of forces causes a moment τ_m_ that induces clockwise rolling as the larva falls back to the surface. For clarity and ease of understanding, the upward movements are exaggerated in panel **C** and **G** of this Figure and the related animations (**Movie S4**). In reality, we’d expect any upward displacement to be very small given the soft nature of the larva. **J)** Spatial trajectories of a soft robotic snake (SRS) attempting to perform rolling in the absence of a solid surface generates a rotational motion that is reminiscent of a semicircular windmill blade rotating around the central post. We refer to this motion as “windmill blade model” (adopted from Arachchige *et al.* with permission). **K)** Experimental validation of the windmill blade model using Drosophila larva. A series of still images demonstrates the “windmill blade” motion of a larva oriented vertically with its tail stuck in an agarose gel, showing the same spatial trajectory as the SRS in J). The screenshots are taken at 0°, 90°, 180°, 225°, and 315° positions during a full 360° “windmill blade” roll. The larva is rolling counterclockwise. The dorsal side of the larva is labeled with a red marker to show the rotation of the body. At 0° position the red mark is invisible but becomes fully visible at 180° position, and becomes invisible again as the larva complete the roll.

Here we describe a discrete and infinitesimal propagation of the bending plane and the subsequent force and moment imbalance to illustrate the larval rolling locomotion. A larva with a flat pose lying on a surface experiences two distinct forces: its weight (*W* = *mg*, where *m* is the mass of the body and g is the gravitational acceleration) and the distributed ground reaction force (F_R_) from the contact points between the body and ground (**Figure 3A**). To achieve initial bending and form a C-shaped body conformation, the larva unilaterally activates (contracts) its ventrolateral muscles (VLs) on the right side (**Figure 3B**). Next, the muscle groups located above the VLs become activated (i.e., clockwise propagation of muscular contraction with respect to the head), causing the head and tail of the larva to undergo an infinitesimal move upward (**Figure 3C**). According to our model, an upward displacement would generate a force imbalance, causing the distributed ground reaction force to act solely on the contact points near the center of the body (**Figure 3C**) as the friction between the larva cuticle and surface (*F_f_*) prevents the body from slipping. Consequently, the reaction force no longer aligns with the body weight (**Figure 3D)**, resulting in a moment of imbalance, *τ*_*m*_, which induces a fall back to the ground and counterclockwise rotation (with respect to the head) to restore the distributed contact (**Figure 3E**). Therefore, during this process, the clockwise progression of muscle contraction causes the larva’s body to undergo a counterclockwise rotation, leading to inward translation of the entire soft body (**Figure 3B-E**). We propose that continuous and successive cycles of imbalance followed by reactive rotation leads to continuous rolling motion. The actual body translation of the larva rotating on the surface results from differential friction reaction forces between the larva’s cuticle (outer coverings) and the contact surface, following the same laws of physics applicable to a tire rotating on a surface. See **Movie S4** for animated version of this model.

On the other hand, if after the initial C-shaped body conformation (**Figure 3F**), contraction of the muscle groups located below the VLs (i.e., counterclockwise propagation of muscular contraction with respect to the head) would infinitesimally move the larva’s mid-body upward while its head and tail are still in touch with the surface (**Figure 3G**). Based on our proposed model, an upward movement would generate a force imbalance that causes the distributed force to act solely on the contact points near head and tail (**Figure 3H**).

Consequently, the reaction force no longer aligns with the body weight, resulting in a moment of imbalance,*τ*_*m*_, which triggers a fall back of the mid-body to the ground, followed by clockwise rotation of the entire body to rectify the statical imbalance and restore stability via distributed contact (**Figure 3I**). Therefore, during this process, the counterclockwise progression of muscle contraction produces clockwise rotation, leading to outward translation of the entire soft body (**Figure 3I**). Based on this model, successive continuous cycles of imbalance followed by reactive rotation lead to continuous rolling motion. See **Movie S4** for animated version of this model.

Our biomechanical models of clockwise and counterclockwise rolling were adopted from two recent robotics studies by Arachchige et al. [39, 40], who extensively modelled the kinematics and physics of rolling and constructed soft robotic snakes (SRS) performing inward and outward rolling motions (**Figure S5A-B and Movie S5**) identical to those of *Drosophila* larvae. The trajectories for these rolling motions were generated from circumferential progression of bending of the entire body, similar to what we observe in larval rolling. Once the SRS is provided with the commands to generate circumferential progression of bending while the SRS is on the ground, it engages in rolling locomotion – both clockwise and counterclockwise directions (the authors termed these gaits are inward and outward rolling) (**Figure S5A-B and Movie S5**) [39, 40].

To further determine the impact of friction reaction forces between SRS skin and the contact surface, Arachchige et al. formulated the mathematical basis of spatial trajectories for an SRS attempting to perform rolling in a three-dimensional (3D) space with no solid surface to interact with its skin. In this simulation, the SRS is anchored to the ground from one end while the rest of its body is up in the air. Based on their formulation, following the initial bending, the SRS forms a C-shaped structure with one end attached to the surface and the other end up in air. Subsequently, while circumferential progression of contraction occurs, the SRS maintains its C-shape and its free end undergoes rotation (circular movement) around the end touching the surface, resembling a semicircular windmill blade rotating around the central post (**Figure 3J**) [39, 40]. This simulated 3D motion (hereafter referred to as “windmill blade” movement) indicated that for circumferentially propagating contraction to be transformed into a rolling behavior, the SRS needs to interact with a surface, thereby generating the friction reaction forces necessary for rolling. We used *Drosophila* larvae to experimentally test and validate the windmill blade movement. We inserted larval posterior segments (A7-A9) into a nick made on an agarose pad, positioning the larva perpendicular to the surface with its head and the rest of the segments free to move in a 3D space (i.e., air) (**Figure 3K and Movie S6**). Upon Goro activation, the larva formed a semicircular C-shaped structure and continuously rotated around its point of contact with agarose (**Figure 3K and Movie S6**). In this setup, if the larva exits the nick and acquires a flat pose lying on the surface, the windmill blade motion is converted back to normal larval rolling. We observed a similar 3D motion in a larva attached to a glass ceiling via only its mouth hook (**data not shown**). In another experiment, we optogenetically activated Goro and first allowed the larva to roll on the surface, and then lifted the animal off the surface while rolling. Once the larva was suspended in the air, it began producing windmill blade movement (**Movie S6**). These data not only corroborate the windmill blade model, but also reveal the role of surface friction in rolling.

Our SCAPE imaging of muscle activity involved activating Goro neurons while the larva was floating in a small chamber filled with water. Due to viscosity the larvae were able to perform escape bending and rolling in-place without translating and without windmilling. The muscle activity we observed during escape rolling, though primarily a circumferential wave of “bend” contractions, also consisted of LT and VA muscle activity that sustained or increased counter to the bend propagation. We propose that these muscles contribute to the mechanics of rolling in two ways. First, it is likely that optimal rigidity must be maintained for efficient rolling of soft-bodied animals. If so, the prolonged activity windows of the larvae’s LT and VA muscles could be involved in adjusting and maintaining internal body pressure, thus ensuring the optimal rigidity necessary for the larval soft body to function as a muscular hydrostat, similar to the hydrostatic skeletons observed in animal tongues and cephalopod arms [41]. Second, transverse (LT) muscle activity might provide a pulling force that aids in generating body rotation. Although our biomechanical model achieves bending and rolling with strictly longitudinal contractile forces, it does not rule out the possible contribution of other muscles with distinct orientations to larval rolling. We test the functional contributions of distinct muscles in the following section.

### The activities of a variety of dorsal, lateral and ventral muscles are required for rolling

Our SCAPE imaging and biomechanical model suggest that a circumferential propagation of muscular contraction underlies the torque required for larval rolling. However, the muscles and MN types that are essential for rolling are unknown. Many larval body muscles are co-innervated by type Is and a single type Ib excitatory MNs **(Figure 4A)**. Each type Is MN innervates multiple target muscles, has a phasic firing pattern, and makes smaller synaptic boutons in neuromuscular junctions (NMJs). Type Ib MNs, by contrast, typically innervate one muscle, have a tonic firing pattern, and establish larger synaptic boutons in their NMJs [42, 43]. There are two phasic firing type Is MNs in each hemisegment, which based on the muscle groups they each innervate, are also known as ventral common exciter (vCE) and dorsal common exciter (dCE) MNs [44]. Here, we sought to understand how the different MNs and muscles contribute to rolling.

**Figure 4:**
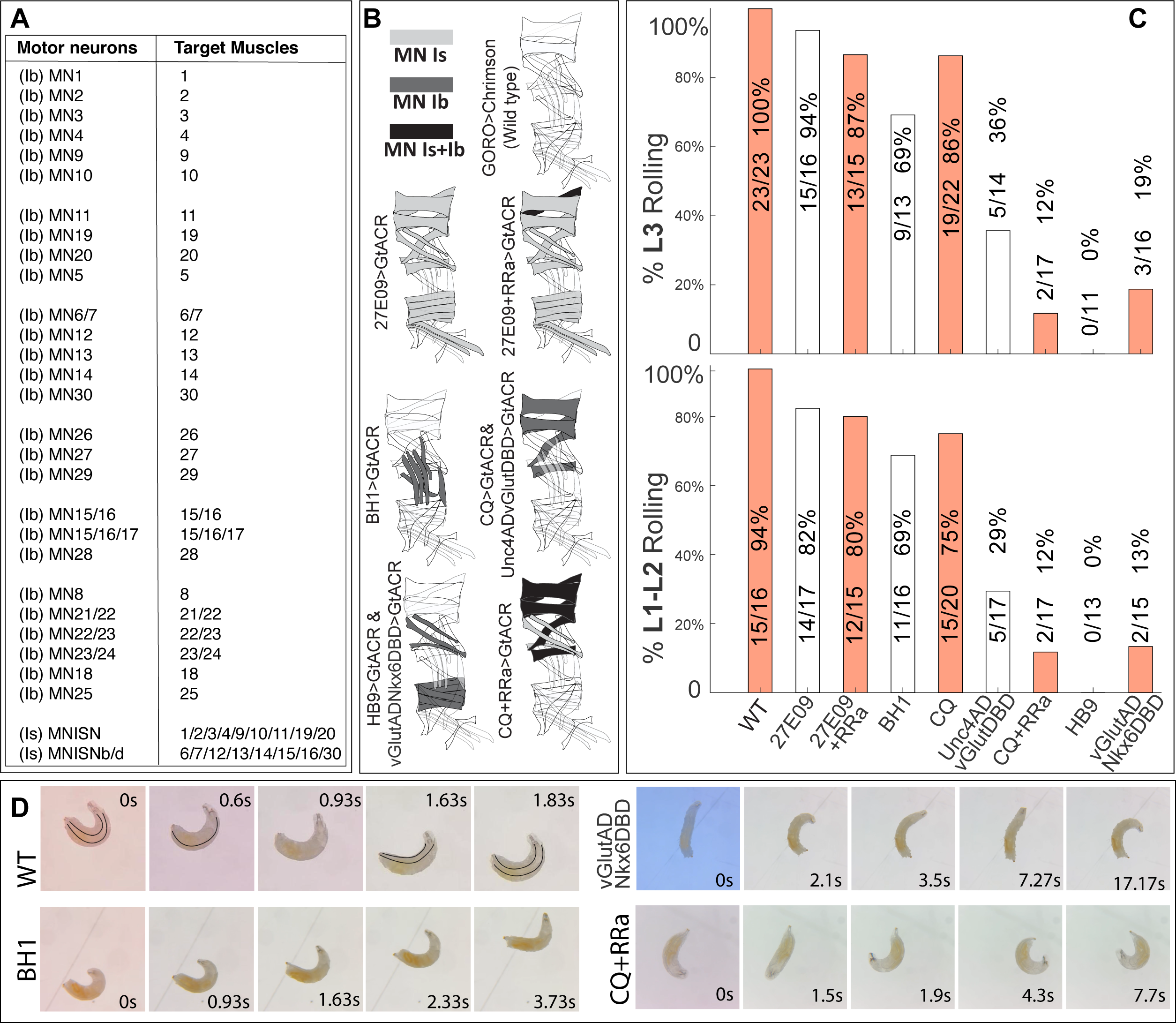
The activity of variety of dorsal, lateral and ventral muscles are required for rolling. **A)** Table listing the larval MN-muscle innervation map. The larva has 27 type Ib MNs each hemisegment. Most of them innervate a single muscle and a few innervate 2 to 3 muscles. There are two type Is MNs each hemisegment (ventral common exciter, vCE and dorsal common exciter, dCE). **B)** Cartoons showing the muscle groups in each genotype whose MN Is (light grey), Ib (dark grey) or both Ib and Is (black) inputs are silenced. *R27E09>UAS-GtACR1-eGFP* silences both MN Is. *R27E09+RRa>UAS-GtACR1-eGFP* silences both MN Is and MN1 (Ib). *BH1>UAS-GtACR1-eGFP* silences Ib MN targeting the lateral transverse (LT) muscles (21,22,23,24,8) and MN5 (Ib). *CQ*>*UAS-GtACR1-eGFP* and *Unc4^AD^-vGlut^DBD^*>*UAS-GtACR1-eGFP* silences Ib MNs of dorsal longitudinal (DL) muscles 2,3,4,9,10. *CQ+RRa> UAS-GtACR1-eGFP* silences the dorsal MN Is in addition to the DL Ib MNs targeted by *CQ. HB9>UAS-GtACR1-eGFP* and *vGlut^AD^-Nkx6^DBD^*>*UAS-GtACR1-eGFP* silences Ib MNs targeting ventral longitudinal (VL) muscles and dorsal oblique (DO) muscles 11,19,20,12,13,6,7,14,30. **C)** - Silencing of different MN groups with *UAS-GtACR1-eGFP* leads to defects in rolling. Bar graphs showing the proportion of L3 (top panel) and L1-L2 (bottom panel) animals able to complete at least one complete roll. Silencing MN Is (*R27E09> UAS-GtACR1-eGFP*) or in combination with Ib MN1 (*R27E09+RRa>UAS-GtACR1-eGFP)* had little or no effect on rolling performance. Silencing Ib MNs of LT muscles (*BH1> UAS-GtACR1-eGFP*) leads to a slightly reduced chance of successful rolling. Silencing Ib MN of DL muscles caused mild (*CQ> UAS-GtACR1-eGFP*) to moderate (*Unc4^AD^-vGlut^DBD^> UAS-GtACR1-eGFP*) defect to rolling; however, silencing both Ib and Is targeting DL muscles (*CQ+RRa> UAS-GtACR1-eGFP*) lead to severe rolling failure. Silencing Ib MNs of VL and DO muscles (HB9> UAS-GtACR1-eGFP and *vGlut^AD^-Nkx6^DBD^*> UAS-GtACR1-eGFP) leads to near complete failure in rolling. Control animals are *69E06-LexA>Aop-Chrimson-tdTomato (“GORO>Chrimson-tdTomato”*), while each MN silencing group also carries one or two *MN-Gal4* or *MN-split-Gal4* components, and *UAS-GtACR1-eGFP*. Numbers on the bars show the number of rolling animals out of the total number of larvae used for each group. **D)** Still images showing wild type (*GORO>Chrimson*) rolling, and rolling in *BH1>UAS-GtACR1-eGFP*, *vGlut^AD^-Nkx6^DBD^>UAS-GtACR1-eGFP* and *CQ+RRa>UAS-GtACR1-eGFP*animals. Wild type rolling is initiated by bending to one lateral side while dorsal plane is facing upward (identified by both tracheae being visible, labeled with black lines), followed by continuous body rotation (still images showing 0°, 90°, 180°, 270° and 360° of an outward roll). A successful roll is defined as the completion of at least one 360° roll. BH1>GtACR (LT Ib MN silencing) animals can roll normally albeit at a slightly slower speed. *vGlut^AD^-Nkx6^DBD^>UAS-GtACR1-eGFP* (VL and DO Ib MN silencing) animals cannot bend their bodies enough to start rolling. *CQ+RRa>UAS-GtACR1-eGFP* (dorsal Ib and Is silencing) animals demonstrate frequently interrupted rolling and inability to maintain rolling in a single direction, where they bend and rotate for about 90°, then bend to the opposite side.

We initially used different MN-Gal4 lines to chronically silence different subsets of MNs with *UAS-Kir2.1-eGFP* (an inwardly rectifying potassium channel [45]), while optogenetically activating Goro command neurons to induce rolling. However, several control experiments led us to question this method. First, we observed weak Kir2.1-eGFP signal in the MNs targeted by *BH1-Gal4* and *CQ-Gal4* throughout development, and lack of expression in some of expected MNs labeled by the *vGlut^AD^-Nkx6^DBD^*split Gal4 line. Moreover, when we examined muscle GCaMP6f activity in intact MN-Gal4>UAS-Kir2.1-eGFP larvae during crawling, we did not detect diminished GCaMP6f signal in the target muscles labeled by *BH1-Gal4*, *CQ>Gal4*, or *CQ+RRa-Gal4* (**Movie S7),** suggesting that the Kir2.1 channel may be ineffective at silencing corresponding MNs. Given the chronic expression of Kir2.1 in these experiments, it is possible that some neurons may have a mechanism to suppress UAS-Kir2.1-eGFP expression or neutralize its effect on membrane potential by changing the array of other membrane ion channels. On the other hand, since MN-Gal4 lines induce Kir2.1 expression in MNs during late embryonic stages, some embryos may be unable to perform the motor behaviors required for hatching and die before emerging as a larva. If so, the successfully hatched larvae may not represent the entire population of animals expressing Kir2.1

Given these caveats, we tested roles for MNs by optogenetically activating Goro command neurons with *Goro-LexA>lexAop-Chrimson*, and acutely silencing subsets of MNs using Gal4-driven GtACR1-GFP, an optogenetically-activated chloride channel [46]. **Figure S6** shows the CNS expression patterns of different MN-Gal4 lines used to drive *UAS-GtACR1-eGFP*. Our dual optogenetic approach allowed us to selectively silence the targeted MNs only during the time window when *Goro-LexA>lexAop-Chrimson* was activated by light, as GtACR1 opens and hyperpolarizes neurons in response to illumination. We used *R27E09-Gal4* to exclusively express *UAS-GtACR1-eGFP* in type Is MNs and found that silencing of these common exciter MNs had no effect on larval rolling. In contrast, optogenetic silencing of type Ib MNs innervating the ventral longitudinal (VL) and dorsal oblique (DO) muscles using *HB9-Gal4>UAS-GtACR1-eGFP* eliminated both bending and rolling in L1/L2 and L3 animals (**Figure 4B and Movie S8**). Given the possible off-target expression of *HB9-Gal4* in other neurons, we repeated this experiment using a recently-developed split Gal4 line (*vGlut^AD^-Nkx6^DBD^*)[47] that specifically targets the same MNs (i.e., VL/DO MNs) as *HB9-Gal4* and observed similar severe rolling and bending defects in L1/L2 and L3 animals (**Figure 4B and Movie S8**). Next, we used *CQ-Gal4>UAS-GtACR1-eGFP* to selectively silence five type Ib MNs innervating the dorsal longitudinal (DL) muscles and found that acute silencing of tonic inputs to DL muscles led to mild rolling defects, where 25% of L1 and 13% of L3 larvae failed to execute at least one complete 360° roll (**Figure 4B and Movie S8**). Since *CQ-Gal4* has been reported to have a few off-targets [48], we repeated the DL silencing experiment using *Unc4^AD^-vGlut^DBD^-Gal4* [49] to specifically express GtACR1-eGFP in the same MNs targeted by *CQ-Gal4* line. *Unc4^AD^-vGlut^DBD^>UAS-GtACR1-eGFP* animals showed 68% and 42% rolling defects in L1/L2 and L3 stages, respectively, which is more severe than what we saw with *CQ-Gal4* (**Figure 4B and Movie S8**). Taken together, the MN silencing data using *Unc4^AD^-vGlut^DBD^* and *CQ-Gal4* lines demonstrate that tonic-firing type Ib MNs innervating DL muscles 2,3,4,9,10 are important for larval rolling. In both *CQ-Gal4> UAS-GtACR1* and *Unc4^AD^-vGlut^DBD^> UAS-GtACR1-eGFP* animals, DL muscles 2,3,4,9,10 still receive phasic excitatory inputs from the type Is MN (dCE) and the DL muscle 1 receives both Ib and Is inputs (**Figure 4B and Movie S8**). The ability of some *CQ-Gal4>UAS-GtACR1-eGFP* and *Unc4^AD^-vGlut^DBD^>UAS-GtACR1-eGFP* animals to perform complete rolls indicates that when Ib inputs to DL muscles 2,3,4,9,10 are blocked, the phasic Is inputs to these muscles, along with the Ib and Is inputs to DL muscle 1, may be sufficient to fully or partially contract the dorsal area of the body wall during larval rolling, thereby preventing any disruption in the circumferential progression of muscle activity required to complete the roll. Therefore, to fully determine the role of DL muscles 1,2,3,4,9,10 in larval rolling, we used *CQ-Gal4+RRa-Gal4* line to silence both Ib (tonic) and Is (phasic) inputs to these muscles. In *CQ-Gal4+RRa-Gal4*>*UAS*-*GtACR1-eGFP* larvae, we saw 88% rolling defect in both L1/L2 and L3 animals, which is significantly higher than the defect seen in *CQ-Gal4+RRa-Gal4>UAS-GtACR1-eGFP* or *CQ-Gal4>UAS-GtACR1-eGFP* larvae (**Figure 4 C-D, Movie S8 and Movie S9)**. Notably, *CQ-Gal4+RRa-Gal4>UAS-GtACR1-eGFP* animals could bend and initiate the roll; however, they could not execute a complete 360° rolling, corroborating that, as suggested in our biomechanical model for rolling (**Figure 3)**, the contraction of DL muscles and the resultant force imbalance are necessary for the larvae to complete the rolling already initiated due to contraction of VL and LT muscles. Finally, we found that silencing Ib MN inputs to lateral transverse muscles (LTs) using *BH1-Gal4>UAS-GtACR1-eGFP* led to a 31% rolling defect, indicating that LTs are in part necessary for rolling behaviors. Given that the LT muscles do not seem to receive any phasic inputs from Is MNs (dCE and vCE), we did not examine larval rolling in *27E09-Gal4+BH1-Gal4>UAS-GtACR1-eGFP*.

To determine the efficiency and specificity of GtACR1-eGFP driven by the Gal4 lines used in this study, in addition to confirming GFP expression in isolated larval brains (**Figure S6**), we performed muscle calcium imaging in intact larvae crawling forward under a confocal microscope. We found that GtACR1-eGFP driven by *vGlut^AD^-Nkx6^DBD^, HB9-Gal4, BH1-Gal4*, and *CQ-Gal4* lines efficiently blocked the activity of the desired MNs, as evidenced by almost complete lack of activity in their target muscles while the larvae performed forward crawling (**Figure S7 and Movie S10**). In contrast, we did not observe diminished GCaMP6f activity in any off-target muscles, indicating the specificity of these Gal4 lines. Owing to the GFP signal from GtACR1-eGFP expression, motor axons were visible in the muscle field, further demonstrating sufficient expression levels of GtACR1-eGFP. In *Unc4^AD^-vGlut^DBD^>UAS-GtACR1-eGFP* animals, although the motor axons clearly expressed GtACR1-eGFP, their target muscles showed low levels of GCaMP6f activity, indicating that they were not fully paralyzed (**Figure S7 and Movie S10**). We reasoned that residual GCaMP6f activity could be driven by a type Is MN (RP2) that provides phasic inputs to multiple dorsolateral muscles, including those innervated by *Unc4-^AD^;vGlut-^DBD^* MNs. To experimentally test this possibility, we performed muscle GCaMP6f activity in *R27E09-Gal4* + *vGlut^AD^-Nkx6^DBD^>UAS-GtACR1-eGFP* animals and found that, consistent with our prediction, Muscles 2,3,4,9,10 showed little or no GCaMP6f activity (**Figure S7 and Movie S10**). Taken together, we conclude that, consistent with our biomechanical model, ventral muscles are essential for initial bending, and muscles located in different dorsoventral regions of the larval body contribute to completing the 360° roll. Furthermore, type Is MNs inputs may act in synergy with type Ib inputs to generate robust rolling behavior and partially compensate for the loss of Ib inputs; while in the absence of Is inputs, type Ib MNs can still drive robust rolls.

### Identifying candidate circuits for circumferential muscle contraction sequences

We next examined electron microscopic (EM) connectome data [21, 50] to determine whether PMN-MN connectivity could provide insights into the basis of bending and rolling. Activity measurements indicated two requirements for the escape motor pattern: 1) left-right asymmetric muscle contraction during bending, and 2) circumferential propagation of this contraction during rolling. To first gain insight into asymmetric muscle contraction, we focused on the spatial distribution of postsynaptic fields on MN dendrites. Previous studies showed that MNs innervating lateral muscles have ipsilateral dendritic arborizations while some MNs innervating midline muscles have both ipsi-and contralateral dendritic arborizations [21, 50–52]. Consistent with this, we found that postsynaptic fields on MNs innervating the lateral muscles are located in the ipsilateral side only. Hence, there is no overlap between the fields of left and right MN counterparts (**Figure S8A,B)**. With this organization, if the larva provides unilateral excitatory or inhibitory inputs to the MNs on one side (e.g., right), the MNs on the other side (e.g., left) will not be affected. Such exclusively unilateral inputs can help larvae perform the initial bending. In contrast, the left and right MNs innervating the dorsal muscles showed a partial overlap in their postsynaptic fields (**Figure S8C**). With this organization, even if the larva sends unilateral excitatory or inhibitory inputs to MNs on one side (e.g., right), the MNs on the other side could still receive and sense some of those inputs. This partial overlap of fields could support the momentary bilateral symmetry that occurs as the dorsal midline rotates into the bend. Such partial overlap was also seen for the left and right MNs innervating the ventral muscles (**Figure S8D,E**), which could support the transient synchronous activity of the left and right ventral muscles while passing through the bend side.

Next, we examined the PMN-MN connectome for circuits that could support circumferential propagation of contraction. First, we identified multiple PMNs that synapse onto MNs with spatially clustered target muscles in the periphery. Specifically, PMNs that synapse primarily to MNs innervating one spatial muscle group are more likely to synapse onto neighboring regions around the circumference of the larva, such as dorsolateral muscles (i.e., DLs, DOs, and LTs), ventrolateral muscles (i.e., VOs, VLs, and LTs) (**Figure 5A, Figure S9A**), and/or muscles flanking dorsal midline (i.e., left-right DLs) (**Figure 5B, Figure S9B**). To verify this phenomenon across all PMNs, we observed the cosine similarity between individual PMN projection patterns relative to downstream muscle drive and found that muscles more proximal along the circumference of the larva have higher overlap in PMN drive than those distal to each other (**Figure S9C**). On the other hand, MNs that innervate spatially distant muscles (i.e., lateral transverse muscles LTs on the left and right side) receive inputs from an exclusive set of PMNs (i.e., right PMNs synapsing with right LTs and vice versa) (**Figure 5C, Figure S9B**).

**Figure 5:**
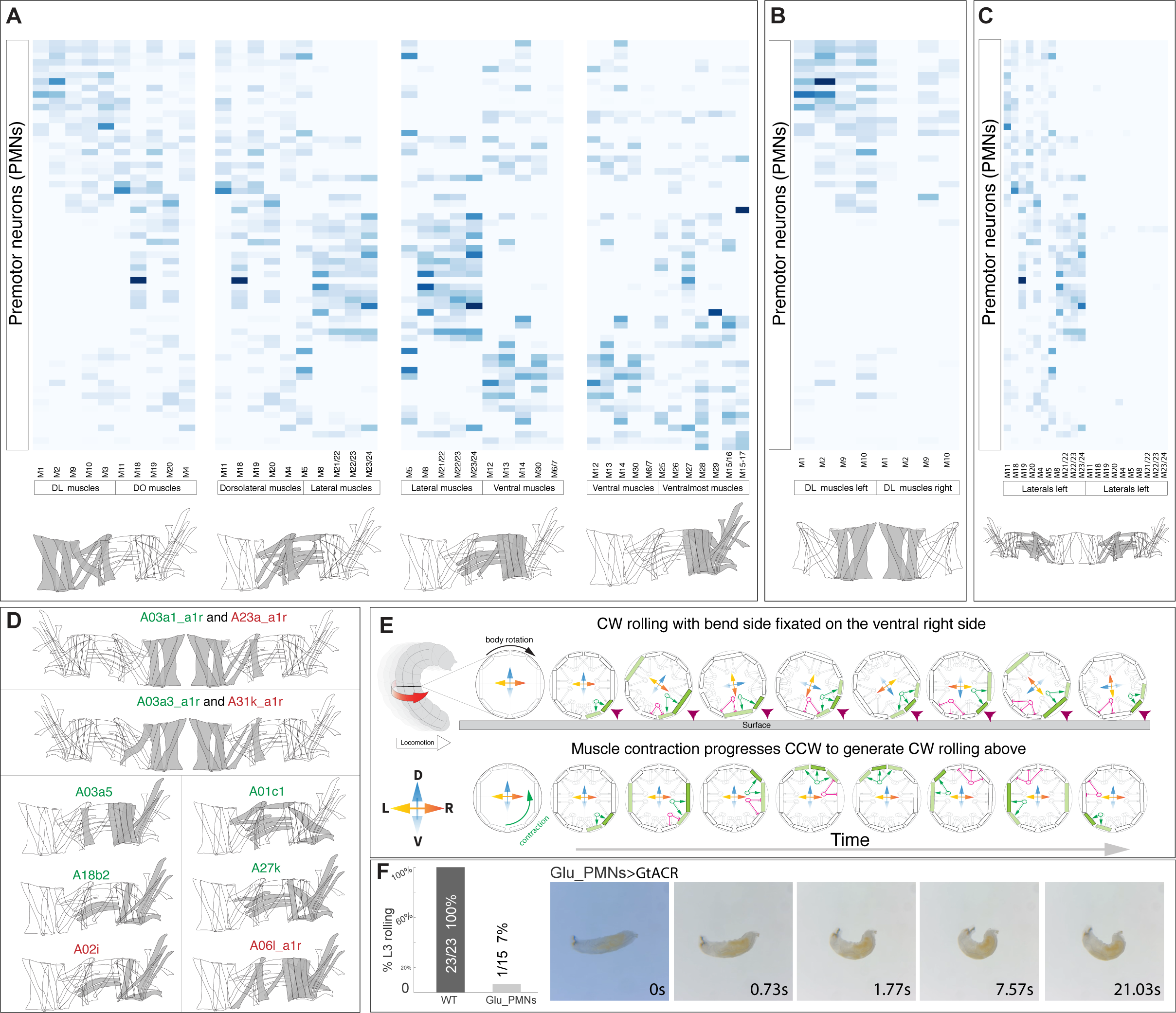
Premotor circuit organization for circumferential progression of muscle activity observed during escape. (**A-C**) Heat maps representing the normalized weighted-synaptic output (blue shading) of left PMN (rows) onto different subsets of left MNs (columns). Grayed sketches at the bottom of heatmaps indicate the target muscles of MNs in heatmap. (**A**) PMNs demonstrate connectivity patterns with dorsoventral organization, where PMNs presynaptic to DL MNs also establish synapses with MNs innervating the neighboring DO muscles (left), PMNs presynaptic to dorsolateral MNs also establish synapses with MNs innervating the neighboring lateral muscles (second from left), PMNs presynaptic to lateral MNs also establish synapses with MNs innervating the neighboring ventral muscles (second from right), and PMNs presynaptic to ventral MNs also establish synapses with MNs innervating the neighboring ventralmost muscles (right). (**B**) Left PMNs presynaptic to left DL MNs also establish significant number of synapses with right DL MNs. Left and right DL muscles span neighboring regions along the dorsal body midline. (**C**) Left PMNs presynaptic to left lateral/dorsolateral MNs have negligible connectivity with right DL MNs. Lateral/dorsolateral on the left and right side are spatially distant from each other. **(D)** Individual PMNs tend to synapse with MNs that correlate with spatially proximal muscles. The target muscles of PMN-MN-Muscle motifs are shown in gray. Excitatory and inhibitory PMNs are shown in green and red, respectively. **(E)** Schematic model showing that to generate clockwise rolling (top panel), muscle contraction progresses counterclockwise (low panel). Circles are cross-section depictions of a larva at different time points. Rectangular shapes around the circumference indicate the body wall muscles. Dark and light green indicate the fully and partially contracted muscles, respectively. Purple arrowheads indicate the fixated bend sides in the top panel. The dorsoventral axis rotates clockwise in the top panel, while it is fixated at the bottom panel. The green and magenta oval shapes indicate the excitatory and inhibitory inputs from active PMNs, respectively. Gray oval shapes indicate inactive PMNs. **(F)** Silencing glutamatergic interneurons severely impairs rolling ability. Bar graph shows that optogenetic silencing glutamatergic interneurons with per>GtACR leads to rolling failure in 93% L3 animals (only 1 of 16 animals rolling) compared to 0% failure rate in control animals (23 of 23 rolling). Still images show that per>GtACR animals attempt to roll by bending but cannot progress to rotate their body in any direction. Control animals are *69E06-LexA>Aop-Chrimson-mCherry (“GORO>Chrimson”)*, while *per>GtACR* also carries *per-Gal4 and UAS-GtACR1-eGFP*.

As a specific example for PMNs driving neighboring muscles, we found that two presumptive excitatory PMNs in the right hemisgment (A03a1_a1r and A03a3_a1r PMNs) are presynaptic to MNs innervating dorsolateral muscles (DLs and DOs) on the animal’s right side, while they also make a smaller number of synapses with MNs innervating DL muscles on the left side (**Figure 5D**). Thus, activity of these right A03 PMNs should strongly activate the right dorsolateral muscles and weakly activate the DL muscles on the left side. Then, activation of the left counterparts of these PMNs should have a mirror effect and strongly activate the left dorsolateral muscles while weakly activating the DL muscles on the right side. Sequential activation patterns of left and right A03 PMNs should therefore facilitate the circumferential progression of dorsolateral muscle contractions from right to left hemisegments, a pattern that is seen during clockwise rolling (**Figure 5E**). Further analysis of connectome data revealed that two inhibitory PMNs (A23a and A31k) have connectivity patterns similar to the excitatory A03a1 and A03a3 PMNs (**Figure 5D**). Such a synaptic organization raises the possibility that circumferential propagation of MN activity could be followed by circumferential inhibition of muscles. As A23a and A31k inhibit dorsolateral muscles, other excitatory PMNs may activate lateral and/or ventrolateral muscles, thereby enforcing wave progression (**Figure 5E**). Consistent with this model, we identified excitatory (e.g., A03a5, A01c1, A18b2, and A27k) and inhibitory PMNs (e.g., A02i and A06l) that synapse with MNs projecting to muscles positioned in the lateral, ventrolateral, and ventral regions of the body wall (**Figure 5D**).

To quantitatively test the significance of the observed circumferential structure of PMN-MN-Muscle connections, we compared the dorsoventral structure of PMN-MN outputs to that which would be expected by chance. Specifically, we performed a shuffling procedure that preserves the general statistics of PMN-MN connectivity while randomizing specific PMN-MN pairs (**Figure S9D,E**). We found that the dorsoventral structure of real PMN-MN-Muscle connectivity patterns of all PMNs combined, as well as only PMNs previously identified as excitatory or inhibitory, is significantly greater than expected by chance (**Figure S9F**). This supports the likelihood that multiple specific premotor motifs could aide in driving a circumferential wave of muscle contractions, like those detailed above, and with precisely timed handoff from excitatory to inhibitory PMNs (**Figure 5D**). Taken together, the PMN-MN-Muscle connectome is structured in way that sequential firing of excitatory and then inhibitory PMNs in one side followed by the activation of their counterparts on the other side could underlie the progression of muscle contraction waves around the circumference of the larva (**Figure 5E**).

There are multiple glutamatergic inhibitory PMNs (i.e., A02e, A02f, A02g, A02h, A02i, A02m, and A02n) that have been implicated in larval crawling and rolling [27, 30, 53, 54]. Based on the PMN-MN-Muscle connectome dataset, each specific A02 PMN establishes synapses with MNs innervating subsets of nearby muscles that together span almost all muscles in the dorsoventral axis of the larval body wall (**Figure S10**). To determine whether glutamatergic PMNs are essential for Goro-triggered rolling, we silenced glutamatergic PMNs using per-Gal4 [53] driven GtACR1-eGFP and quantified the rolling success rate. We found that optogenetic silencing of per-Gal4 neurons led to significant decreases in the rolling success rate in the L3 animals (**Figure 5F and Movie S11**). It is likely that in addition to PMNs, per-Gal4 targets other glutamatergic interneurons. Therefore, it will be of great interest to functionally test and determine the role of individual or smaller subsets of glutamatergic PMNs as well as other excitatory and inhibitory PMNs in generating rolling behavior.

## Discussion

Escape is a fundamental form of locomotion and critical for the survival of all animals. To understand the neural mechanisms of an escape behavior, we have performed live imaging of muscle activity across animals as they perform rolling escape behavior, analyzed a connectome for motor circuits that could support the unique muscle propagation wave that coordinates this escape motor program, and performed MN silencing experiments to determine muscle groups whose activity is necessary for rolling behavior. This work has illuminated fundamental distinctions and similarities between motor patterns underlying forward crawling and escape in the larva, and starts to uncover the circuit basis for rolling escape motor response.

### Enhancements of SCAPE microscopy to permit studies of rolling behavior

Our ability to identify patterns of muscle activity during rolling was aided by the further development of dual-color SCAPE. High-speed, high-resolution SCAPE microscopy has previously captured muscles in behaving larvae [31] and dual-color proprioceptor activity during crawling behavior [33]. Expanding dual-color SCAPE imaging here enabled ratiometric quantification of muscle activity and discrimination of activity signals from passive changes in fluorophore density within muscles. This imaging method also allowed measurement across muscles at different focal depths during freely moving behavior, including muscles within the C-shaped bend during rolling escape behavior. We expect that further development of SCAPE, for example wider field of view optics and simultaneous collection of signals from multiple angles, will allow coincident imaging in the CNS and muscles and test some of the hypotheses about PMN to MN transformations that underlie rolling, and a number of other behaviors, in simple model organisms.

### Separable sequences of muscle activity during rolling

Animal movement universally involves coordinated sequential muscle activity. We find that the segmentally synchronous circumferential progression of muscle contractions define escape rolling behavior in *Drosophila* larvae. This sequence of activity progresses in a clockwise or counterclockwise manner, which determines the direction of rolling. A recent independent investigation of the motor pattern that generates larval rolling escape behavior similarly found that a circumferential wave of muscle activity occurs during rolling, and that the sequence of this wave determines the rotational direction of the larva [55]. Whether the circumferential sequence is the primary motor activity that promotes rolling, or whether other independent patterns are involved, is so far unclear. However, our muscle imaging revealed evidence that, during rolling, LT and VA muscles contract as part of a separate, out-of-phase sequence relative to the major wave of activity of longitudinal muscles. LT muscles begin to contract after D and V longitudinal muscles also during crawling [21, 50, 54], suggesting special roles for these muscles in both forms of locomotion. During forward crawling, contraction of LT muscles shortens the dorsoventral axis, providing a force that is thought to push the neighboring internal organs to the anterior side [20]. We propose that during rolling, sustained LT and VA muscle activity ensures the optimal rigidity necessary for the larval soft body to function as a muscular hydrostat, which is reminiscent of the structural rigidity required for function of boneless hydrostatic skeletons, such as animal tongues and cephalopod arms [41]. Interestingly, while silencing the LT-innervating MNs substantially compromises rolling behavior, it does not seem to perturb the execution of forward peristalsis itself [27], but reduces the peristalsis frequency and thereby crawling speed [56].

One crucial future goal will be to understand how segmentally synchronous contractions of specific groups of muscles are initiated and then how contraction propagates to other muscle groups in a precise circumferential sequence during rolling. The fundamental distinctions between bodily coordination during rolling versus crawling illustrate the remarkable propensity for even relatively simple nervous systems to generate vastly different circuit dynamics based on context. Prior to this work, it was unknown whether rolling escape would involve a peristaltic wave component, like that observed in crawling, and our work argues against a peristaltic component to rolling. Understanding at a circuit level how the larval motor system switches between these two patterns of activity remains an important future goal. Despite these global differences, the analogies in LT activity in muscle contraction sequences during crawling and rolling raise the question of how translatable our knowledge of premotor crawling circuits will be for understanding rolling. It will be especially important to determine whether excitatory and inhibitory microcircuit motifs that coordinate crawling [43, 57] also coordinate rolling muscle contractions. Further, understanding how the larva transitions between peristaltic and rolling escape locomotor modes remains an open question.

### Laws of mechanics underlying larval rolling escape behavior

We present biomechanical models aiming to explain how counterclockwise circumferentially progressing muscular contraction generates clockwise rolling motion, and vice versa. In our proposed models, which were adapted from the works of Arachchige et al. [39, 40], the rolling behavior is a stabilizing response generated via instantaneous torque (moment) imbalance owing to the offset of the ground reaction forces and weight (i.e., center of gravity, CoG). In a straight larva lying flat on a surface, the CoG is located at the center of the anteroposterior axis of the body (centroid), and the weight cancels the ground reaction forces. For our rolling biomechanical model to work, the larva needs to bend to the either-side to form a C-shape, causing the CoG to move away from the central axis. We expect that for the most efficient rolling escape maneuver, there should be an optimal C shape, and the larval body with either of the two extreme shapes, letter I-shape and letter O-shape, should fail to produce translation on the surface (i.e., movement). A segmentally repeated pair of interneurons, known as down and back (DnB), has been shown to play a key role in bending and rolling [30]. Animals with silenced DnB neurons can perform rolling but with a significantly reduced rolling frequency. Importantly, silencing DnB neurons dramatically decreased the curvature of the C-shaped bending [30]. It would be of great interest to determine if the reduced curvature resulting from DnB silencing compromises the distance that the larva travels (translates) per roll. Given that DnB neurons receive direct inputs from sensory neurons responding to noxious stimuli and gentle touch, it will also be interesting to determine if there is any correlation between the level of noxious input, curvature of the C bend, and the translation distance travelled per roll. Optogenetically activating nociceptive neurons with different intensities of light could be a possible way to regulate the intensity of the noxious insult. DnB neurons are cholinergic excitatory neurons that provide direct synaptic input to multiple excitatory and inhibitory premotor neurons. Selective silencing of these premotor neurons is yet another exciting future direction aimed at understanding how the coordinated activity of different groups of excitatory and inhibitory premotor neurons results in the optimal curvature and circumferential propagating muscular activity essential for roll generation.

### Insights into motor circuits critical for rolling behavior

Our functional manipulations of MNs innervating distinct muscle subgroups highlight the modularity of motor control in larval rolling escape behavior. Larvae can exclusively bend, or perform bending and rolling. These behavioral components occur probabilistically and are dependent on the precise nature of sensory input to the larva [24, 26, 30]. Some sensory interneurons are necessary and sufficient for escape bending to occur but are not necessary for escape rolling [30]. Namely, we see that a circumferential wave of bend-promoting contractions occurs in longitudinal muscles, while counteracting contractions occur in LT and VA muscles that could provide a pulling force and/or regulate body rigidity. Our MN silencing experiments demonstrate that longitudinally-spanning ventral muscles are essential for larval bending and rolling altogether, while longitudinally-spanning dorsal muscles and lateral transverse muscles are only essential for completion of rolling. These findings are congruent with previous work demonstrating that neurons driving transverse muscles are essential for body rotation in self-righting [58] and rolling [27]. Uncovering the premotor circuit modules that drive the bend and the roll could provide insight into how the larval nervous system generates many motor components of its behavioral repertoire.

The relative roles of Is and Ib MNs in different larval behaviors is still debated [42, 59–61]. Our MN silencing experiments showed that silencing type Ib MNs led to more severe rolling defects in rolling behavior than when the broadly projecting type Is MNs were silenced. Furthermore, the simultaneous silencing of Ib and Is MNs led to more serious rolling defects than when only Ib MNs were silenced, suggesting that phasic Is inputs may partially compensate for the loss of tonic Ib inputs. Phasic Is MNs have a higher probability of vesicle release at the NMJ, demonstrate elevated presynaptic calcium influxes upon stimulation, and contain larger synaptic vesicles per synapse than tonic Ib MNs, leading to higher amplitude EPSPs in muscles following type Is activity [59, 62, 63]. For these reasons, Ib and Is MNs are sometimes called “weak” and “strong” MNs, respectively [59, 62]. On the other hand, type Ib NMJs have higher levels of readily-releasable vesicle pool, a higher number of active release sites than Is boutons, and are recruited earlier and for a longer duration during larval movement [59–62]. These physiological properties of type Ib tonic and type Is phasic MNs are determined by their distinct transcriptional profiles [63], and permit a division of labor during muscle contraction, where type Ib activity can coordinate specific, finer muscle contraction timing with varying levels of contractile force upon low-level premotor input, while type Is activity is recruited at high-level premotor input to increase contractile force of ongoing behavior or make large, forceful shifts in movement [59–62]. In conjunction with our results, these properties suggest that type Ib MNs are crucial for driving the main muscle contraction pattern underlying rolling, while type Is MNs merely contribute additional contractile force to rolling, consistent with the minimal impact of type Is MN silencing on crawling behavior [60]. These findings are also consistent with tonic and phasic motor control principles observed in mammalian systems [64–66], adult *Drosophila* walking [67] and flight [68], and other escape motor circuits [18].

We uncovered MN post-synapse distributions and patterns of PMN-MN connectivity that could support propagation of muscle contractions around the circumference of the larva during rolling behavior. Studies of PMNs that innervate midline muscles could reveal how the circumferential wave progresses from left to right sides of the larva’s body. Specifically, PMNs that demonstrate bilateral drive to midline muscles essential for bending and rolling could be necessary for action selection and maintenance of specific escape rolling directions by permitting the circumferential sequence across left and right boundaries of the body. Bridging the gap between Goro escape-promoting command neurons to specific excitatory and inhibitory PMNs through further connectome reconstruction and circuit manipulation is a crucial next step in understanding the basis of larval rolling escape behavior. Continued pursuit of the sensorimotor circuits that drive escape in the larva will more broadly uncover the circuit mechanisms responsible for transforming sensory information into robust and flexible behaviors across taxa.

## Materials and Methods

### Key Resources Table

| Reagent type (species) or resource | Designation | Source or reference | Identifier s | Additional information |
| --- | --- | --- | --- | --- |
| Genetic reagent (*Drosophila melanogaster*) | w1118 | BDSC |  | Global heat + vibration assay line, crossed as control for MN silencing experiments |
| Genetic reagent (*Drosophila melanogaster*) | R69F06-Gal4 | BDSC | # 39497 | Goro line |
| Genetic reagent (*Drosophila melanogaster*) | 44H10-LexA, Aop-GCaMP6f; Aop-mcherry, UAS-Chrimson-mcherry | Aref Zarin |  | Optogenetics line |
| Genetic reagent (*Drosophila melanogaster*) | 44H10-LexA, Aop-mcherry; 44H10::GCaMP6f, UAS-Chrimson-mcherry | Aref Zarin |  | SCAPE ratiometric muscle imaging |
| Genetic reagent (*Drosophila melanogaster*) | 69E06-LexA; Aop-Chrimson-mCherry, UAS-Kir2.1-eGFP | Aref Zarin |  | Goro activation and MN silencing |
| Genetic reagent (*Drosophila melanogaster*) | 69E06-LexA; Aop-Chrimson tdTomato, UAS-GtACR1-eGFP | Aref Zarin |  | Goro activation and MN silencing |
| Genetic reagent (*Drosophila melanogaster*) | 27E09-Gal4 | BDSC |  | MN silencing |
| Genetic reagent (*Drosophila melanogaster*) | RRa-Gal4 | Albert Cardona |  | MN silencing |
| Genetic reagent (*Drosophila melanogaster*) | BarH1-Gal4 (w[*]Pw[+mW.hs]=GawB NP4099 / FM7c) | Kyoto Stock Center | DGRC# 104-601 | MN silencing |
| Genetic reagent (*Drosophila melanogaster*) | CQ-Gal4 | Miki Fujoka, Jim Jaynes |  | MN silencing |
| Genetic reagent<br>(*Drosophila melanogaster*) | *Unc4<sup>AD</sup>-vGlut<sup>DBD</sup> split Gal4 line* | Gift from Haluc Lacin |  | MN silencing |
| Genetic reagent<br>(*Drosophila melanogaster*) | *vGlut<sup>AD</sup>-Nkx6<sup>DBD</sup> split-Gal4* | Gift from Haluc Lacin |  | MN silencing |
| Genetic reagent<br>(*Drosophila melanogaster*) | 44H10- UAS-Kir2.1-EGFP; GCaMP6f | Aref Zarin |  | MN silencing along with muscle calcium imaging |
| Genetic reagent<br>(*Drosophila melanogaster*) | 44H10-GCaMPf, UAS-GtACR1 | Aref Zarin |  | MN silencing along with muscle calcium imaging |

### Contact for Reagent and Resource Sharing

Further information and requests for resources should be directed to and will be fulfilled by the Lead Contact, Aref Zarin.

### Experimental Model and Subject Details

*Drosophila* melanogaster strains were reared on standard molasses food (agar, cornmeal, yeast, molasses, methylparaben, ethanol, propionic acid, penicillin, streptomycin) at 25°C, 60% humidity. Animals of either sex were used. For behavior experiments, third instar larvae were used. For muscle imaging experiments, late first and early second instar larvae were used.

### Method Details

#### Behavior Experiments Using FIM Table

For behavior experiments, at least two sessions, performed on separate days, were performed for each genotype. Larvae were only tested once. Sample sizes for global heat + vibration and optogenetic Goro activation escape assays were designed to replicate recently published larval escape assay data^26^. Rolling escape pattern frequency and initial kinematic characterization experiments were conducted using the FIM (Frustrated total internal reflection Imaging Method)^59^. Videos were acquired at 50 frames per second with a Basler ACE acA2040-90uc four megapixel near infrared sensitivity enhanced camera equipped with CMOSIS CMV4000 CMOS sensor. Camera was equipped with LM16HC-SW lens (Kowa), and BN880-35.5 filter (Visionlighttech). IR diodes (875 nm, Conrad) were used for FTIR imaging and images were acquired using Pylon camera software (Basler). Animals were placed on 0.8% agar surface ∼2 mm thick (Molecular grade, Fisher Scientific).

For the global heat and vibration assay, we developed a novel implementation of FIM, where the agar surface was uniformly heated and temperature was read out using a Elitech STC-1000 and two 10 kOhm temperature probes. These probes measured temperature in the FIM table chamber, which was heated by four LUBAN Mini Hot Air Guns, and the agar substrate itself. Third instar w1118 larvae were rinsed and transferred to a petri dish with 0.8% agar at least 10 minutes prior to experimentation. Then, larvae were gently transferred with a paintbrush to the agar surface. Vibration was generated using Logitech Multimedia Speakers Z200 with Stereo Sound for Multiple Devices. The composite 1000Hz and wasp sound, played at 100dB, was previously published in Wreden *et al.*, 2017^60^. Behavior was recorded immediately.

For optogenetic Goro activation experiments, larvae were raised in the dark on molasses food with 1 mM all-*trans*-retinal (ATR). These larvae were offspring of *44H10-LexA, Aop-mcherry; 44H10::GCaMP6f,UAS-Chrimson-mcherry* female and *R69F06-Gal4* male flies. Larval care prior to experimentation was the same as in the global heat assay, above. Two rings of Blue (470 nm) LED lights (WFLS-G30 × 3 WHT, SuperBright LEDs) of 5 inches and 8 inches diameter were placed approximately 5 inches underneath the FIM table. LED brightness was programmed for experiments using custom code in ARDUINO, as the LEDs’ pulse width modulation was controlled by an ARDUINO Mega 2560 board. For Goro activation experiments, blue LEDs were active at 2230 uW/mm^2^. Larvae that failed to move at all during trials were excluded from any analysis. Bend and roll pattern frequencies were quantified manually by evaluating TIFF stacks in FIJI (https://imagej.net/Fiji).

#### Neural Silencing Combined with Rolling Assays

To assess the roles of different MNs or PMNs on rolling escape, we performed functional silencing of MN-Gal4 lines. Rolling was induced and imaged in larvae carrying the LexA driver for Goro neurons and Chrimson-mCherry. Constitutive silencing of MNs was achieved via the expression of the inward-rectifying potassium channel, Kir. *69E06-LexA; Aop-Chrimson-mCherry, UAS-Kir2.1-eGFP* females with MN-Gal4 males were crossed for experimental groups, and *69E06-LexA; Aop-Chrimson-mCherry, UAS-Kir2.1-eGFP* males with w1118 females were crossed for control groups. For the experiments involving CQ-Gal4 and BH1-Gal4 driven Kir2.1-eGFP expression, L1 larvae were used due to reduced transgene expression in later instars. For all other Kir2.1 crosses, L3 larvae of the experimental and control groups were used. Optogenetic silencing of MNs was achieved via the expression of an anion channelrhodopsin, GtACR1-eGFP*. 69E06-LexA; Aop-Chrimson-tdTomato, UAS-GtACR1-eGFP* flies were crossed with *MN-Gal4* or w1118 lines for experimental and control groups, respectively. For all crosses involving MN silencing with GtACR1-eGFP, L1, L2, and L3 larvae of the experimental and control groups were used. For muscle calcium imaging experiments, we used GMR44H10-GCaMP6f transgenic fly to selectively express GCaMP6f in the body wall muscles [21]. To determine if MN-Gal4 driven expression of GtACR1-eGFP or Kir2.1-eGFP silences the activity of target muscles, we crossed 44H10-UAS-Kir2.1-EGFP; GCaMP6f or 44H10-GCaMPf, UAS-GtACR1 to the different MN-Gal4 lines and performed muscle calcium imaging in offspring larvae.

For both constitutive and optogenetic silencing experiments, larvae were raised on molasses food in dark and fed with 1mM all-trans-retinal (ATR) 24 h prior to the experiment. Larvae were picked from food, rinsed, and given a 30-second rest before imaging to avoid any effect of the minimum light exposure during the process. The animals were then placed onto a 1.5% agarose pad in a 10cm round Petri dish and imaged with an iPhone SE (second generation, Apple Inc.) camera at 30fps. For L1 animals, the larvae and agarose gel were placed under a Stemi 508 dissection scope (Zeiss) and imaged by the iPhone SE camera through a microscope phone adapter (Gosky) with 16X built-in eyepiece (the original 10X eyepiece was removed from the dissection scope). Optogenetic activation of rolling was induced with 30 seconds of intense, uniformly applied white light from beneath the agarose gel (Oeegoo 18W square LED ceiling light) after a 10-second dark period. To ensure the expression of GtACR1-eGFP and Kir2.1-eGFP in the neurons of interest, following the completion of the behavioral assays, animals were dissected and GFP expression was confirmed in isolated CNS using a confocal microscope (Zeiss LSM 900).

Rolling performance was scored manually after blinding data. Rolling success was defined as a larva completing at least one 360° roll (after body bending, able to rotate in a single direction from dorsal side up to dorsal side up). Video processing was done with freeware FFmpegtool (https://ffmpeg.org/) and ImageJ (https://imagej.net/, National Institutes of Health, Bethesda, Maryland).

#### Confocal Image Acquisition

Muscle activity during crawling was initially measured in Zarin *et al.*, 2019 [21]. Briefly, images were acquired using a 10x objective on an upright Zeiss LSM800 microscope. GCaMP6f and mCherry fluorescence values were extracted from a previously acquired forward crawling video and ratiometric values from muscle ROIs (see SCAPE analysis) were subsequently compared to ratiometric muscle ROI values from SCAPE escape images.

#### SCAPE Image Acquisition

A custom-built Swept Confocally Aligned Planar Excitation (SCAPE) microscope, extended from that described in Voleti *et al.*, 2019 [35] & Vaadia *et al.*, 2019 [33], was used to acquire high-speed volumetric imaging of rolling larvae. The system utilizes a custom-made water chamber between the second and third objective lens to accommodate higher collection efficiency over a large field of view with 1.0 NA water immersion objectives throughout. 488nm and 561nm lasers were used to excite GCaMP and mCherry simultaneously, while also activating Chrimson for optogenetics. An image splitter with 561nm long pass dichroic mirror was used with 525/50 and 618/50 filters to split the image into two sCMOS cameras (Andor Zyla 4.2 plus) which enables acquisition of both green and red channels simultaneously with a field of view larger than 1.1mm in the lateral dimension. A 3D field of view with the size of the agar arena was acquired at 10 volumes per second to minimize the motion blur from the larva rolling behavior.

Larvae were raised and fed ATR for optogenetic activation as mentioned above. Larvae were imaged while in a thin 2% agarose chamber filled with distilled water. Each larva was assessed for brightness of muscle fluorophore expression and likelihood of rolling through epifluorescent screening, and the best overall larvae were placed into agar chambers using a fine paint brush. Each chamber was made using 3D-printed molds of different dimensions for late L1 and early L2 larvae of slightly different sizes.

#### SCAPE Image Processing & Analysis

SCAPE volumetric data are inherently three-dimensional. To overcome 3D-segmentation difficulties but keep the richness of volumetric muscle data, we developed a custom MATLAB script to generate a 3D-averaged projection of all SCAPE data. Our 3D-averaging method generates separate GCaMP and mCherry 2D images dynamically based on fluorescence histograms of the z-depth at each x,y pixel location, allowing us to isolate and average signal exclusively from muscle tissue. 3D-averaged projection images were then analyzed using modified versions of the custom MATLAB script from Zarin *et al.*, 2019 and a custom MATLAB script for measuring ratiometric signal and muscle length rapidly using interactive linearly interpolated linescans. For ratiometric signal vs. length measurements, lines were drawn on select frames for example muscles segments A1-A6 on two rolling SCAPE ratiometric movies, and linear interpolations of linescans on intermediate frames were checked frame-by-frame for each movie. Ratio values and muscle lengths were quantified using custom MATLAB scripts. For ROI-based rolling muscle activity analysis, ROIs were drawn manually on non-overlapping portions of each muscle for each segment from A2-A4 on a frame-by-frame basis. Because puncta were present throughout muscles in the mCherry images, pixels with a fluorescence value greater than 90% of the distribution of fluorescence values within the mCherry ROI were discarded in both the mCherry and GCaMP channels to prevent biased ratiometric signal. The ratio of fluorescence between the ROI pixels of the mean depth-projected GCaMP image and the mean-depth projected mCherry image was calculated. The mean of this ratiometric value was extracted for each time point of the ROI, generating individual muscle activity traces. Four roll bouts (one complete revolution based on muscle locations) from four larvae were selected for rolling muscle activity analysis. Subsequent analyses were performed using custom MATLAB scripts.

For comparisons between ratiometric muscle activity values of crawling and rolling, normalized time at maximum ratiometric values were extracted. Statistical analyses were performed as described below. For bend vs. stretch side comparisons of individual muscle activity during rolling, ROI centroids were tracked and, according to the bend and rotation directions of the larvae, divided into four quartiles based on spatial position within the larva’s body at each time point. Ratiometric values from the most bent quartile and ratiometric values from the most stretched quartile were extracted and averaged for each muscle. These values were then compiled, z-scored, and averaged across segments and across three roll videos, providing a “bend” and “stretch” ratiometric values for each muscle, shown in the boxplots in **Figure 2A**. Statistical analyses were performed using custom programs written in MATLAB. For muscle activity peak differences, Wilcoxon Rank Sum tests were used.

#### Electron microscopy, CATMAID reconstructions, and Quantitative Analysis

PMN-MN connectome previously reconstructed in Zarin *et al.*, 2019 were used to generate **Figure 5, Figure S8, Figure S9, and Figure S10.** For quantitative analysis of the dorsoventral structure of PMN-MN connectivity, each MN was assigned a spatial position 1-30 to reflect how their projections map onto the dorsoventral axis of muscles in the body wall. For example, DL muscles 1, 9, 2 and 10 were assigned position numbers 1, 2, 3, and 4, respectively; while VO muscles 28, 15, 16, and 17 were assigned position numbers 27, 28, 29, and 30, respectively. A weighted average was calculated for each PMN using this MN position assignment. PMNs were then sorted by their strength of connectivity to dorsoventrally organized MNs, and averaged left and right PMN-MN weights were z-scored and displayed (**Figure S9A**). Following this sorting of PMNs based on their spatial connectivity strengths, the connection weights of PMNs to MNs were binarized (**Figure S9D**). Cosine similarity analysis was then performed between each PMN, providing a metric of similarity, or more literally, distance between the pairwise binary vectors of each PMN’s spatially organized outputs. A cosine similarity of 1 means pairwise PMN-MN connectivity vectors overlap perfectly, or have identical outputs, while a cosine similarity of 0 means pairwise PMN-MN connectivity vectors are orthogonal, or have no common outputs (**Figure S9C**).

To determine whether global PMN-MN connectivity patterns demonstrate greater dorsoventral patterning with respect to downstream muscles than expected by chance, a shuffling procedure was employed, similar to that used in Hayashi *et al.*, 2022 [69]. Specifically, to generate a statistically meaningful null hypothesis of how PMN-MN connectivity could be structured, binary weights from each PMN onto all MNs were randomly selected with the number of outputs that each PMN makes and the probability of inputs to each MN constrained by the properties of the real PMN-MN connectivity matrix. This constrained global statistics of the 1000 shuffled connectivity matrices to match the statistical properties of the real connectivity matrix without constraining specific PMN-MN partner identities (**Figure S9E**). The relative dorsoventral structure of real PMN-MN connectivity compared to null connectivity was assessed by performing linear regression on the weighted averages of real vs. shuffled PMN outputs and comparing the regression coefficients between real and shuffled data (**Figure S9F**). The regression coefficients serve as a metric of how strongly PMNs prefer MNs based on muscle dorsoventral order, and the regression coefficients for real connectivity data lie outside the 95% confidence interval of the null distribution of regression slopes, demonstrating statistical significance. Unilateral vs. bilateral PMN-MN projection comparisons according to spatial muscle groups were performed by calculating the weighted average of bilateral vs. unilateral inputs onto MNs, and statistical difference between pairs was determined using the Mann-Whitney U test (**Figure S9B)**.

## Data and Software Availability

Behavioral analysis, 3D averaging, signal extraction, and subsequent muscle activity analysis were performed using custom MATLAB scripts that have been included with manuscript submission and are also available here: https://github.com/cooneypc4/larval_escape_manuscript.

Larva 3D model and animation were generated with Blender 3.3.9 (Blender Foundation). Figures and supplementary videos are assembled with Adobe Illustrator 2023 and Adobe Premiere Pro 2023 (Adobe).

## Supporting information

## Acknowledgements

We thank Ashok Litwin-Kumar for providing guidance on quantitative analysis of connectome data. We thank Albert Cardona, Richard D. Fetter, and the HHMI Janelia Fly EM Project Team for providing the raw data of the whole CNS EM volume. We thank Akira Fushiki and Maarten Zwart for annotating neurons, and Keiko Hirono for generating transgenic constructs. We thank Grace Shin for assisting with behavioral assay development. We thank Malte Casper for supporting SCAPE data processing. Stocks obtained from the Bloomington Drosophila Stock Center (NIH P40OD018537) were used in this study. Research reported in this publication was supported by an institutional startup fund from Texas A&M University (AAZ), NIH NINDS R01NS061908 (W.B.G), NIH BRAIN 5U01NS094296 and 1UF1NS108213 (E.M.C.H. and W.B.G.), DoD, MURI W911NF-12-1-0594, and Simons Foundation Collaboration on the Global Brain (E.M.C.H.), and an NSF predoctoral fellowship DGE 2036197 (PCC). The content is solely the responsibility of the authors and does not necessarily represent the official views of the National Institutes of Health.

