## Supplementary material for "Neuromuscular Basis of *Drosophila* Larval Rolling Escape Behavior"

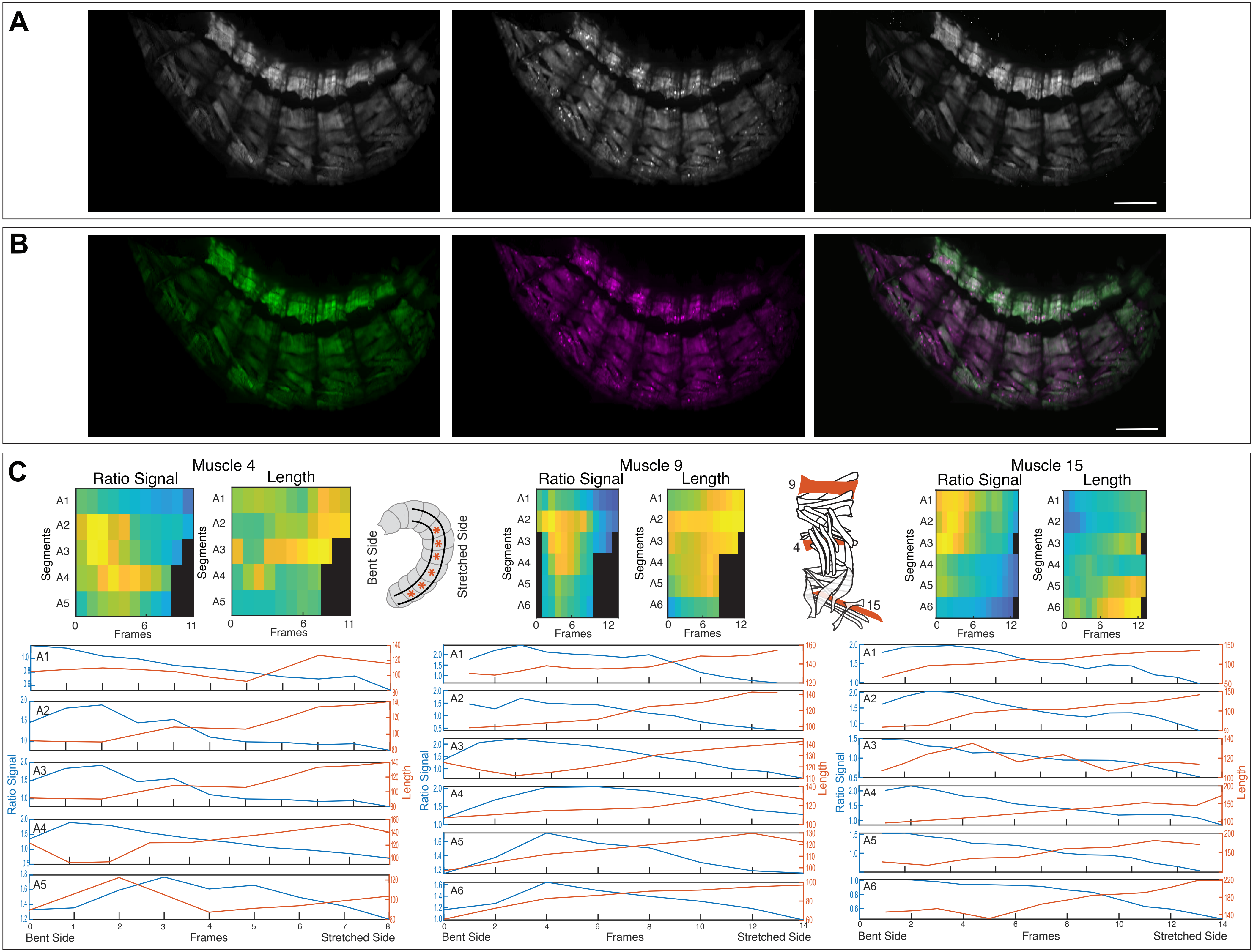
A
B
C
Muscle 4
Ratio Signal
Length
Segments
A1
A2
A3
A4
A5
Frames
Bent Side
Stretched Side
Muscle 9
A6
9
4
15
Muscle 15
Ratio Signal
Length
Bent Side
Frames
Stretched Side

**Figure S1: Dual-color SCAPE permits quantitation of individual muscle activities across larval segments during escape.** (**A,B**) Example stills from GCaMP, mCherry, and calculated ratiometric SCAPE images in grayscale (**A**) and in respective green and magenta and merged (**B**). (**C**) Heatmaps of ratiometric signal and length measurements extracted from example muscles 4 (left), 9 (middle), and 15 (right) across segments A1-A6 during escape rolling. Schematics indicate segments measured and example muscles measured. Below, ratiometric signals (blue) and length measurements (orange) extracted from example muscles during rolling. Plots demonstrate inverse relationship between ratiometric signal and muscle length, demonstrating that ratiometric signal serves as adequate quantitation of muscle contraction.

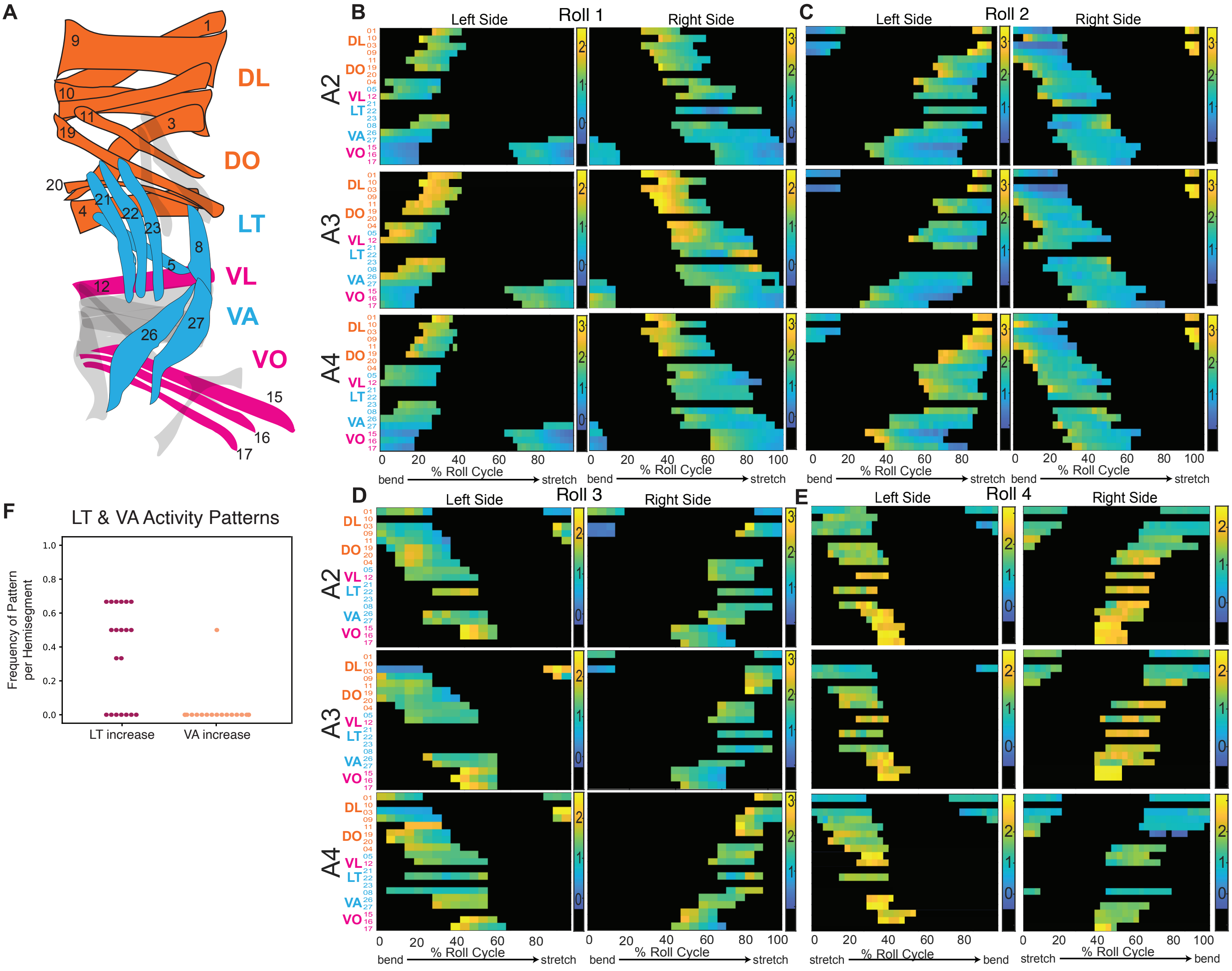


**Figure S2: Individual muscle activity patterns show symmetric dorsoventral propagation on left and right sides. (A)** Hemisegment color-coded by muscle groups that correspond with color-coded muscle labels to the left of the rows in heatmaps. These color groupings match the population bend vs. stretch fluorescence comparisons in Figure 2A. **(B-E)** Heatmaps of z-scored ratiometric signal across all measured muscles, divided by segment (A2-A4) and side (left and right). Each row for each heatmap panel is an individual muscle, organized from dorsal (top row) to ventral (bottom row), and color-coded according to muscle groupings in A. Colorbars to the right of the upper heatmaps show range of ratiometric values. **(B-D)** Traces from muscles moving out of the bend during rolling show moderate decrease in ratiometric signal. **(E)** Muscles moving toward the bend show mild increase in ratiometric signal. Notably, ratiometric signals in muscles moving into the bend from our imaging perspective have a less apparent activity pattern. **(F)** Frequency of muscles within each analyzed hemisegment within LT and VA groups that demonstrate relative increases in ratiometric signal as they rotate toward the bend, where 0 indicates that no muscles in that hemisegment demonstrated increased activity in the bend phase and 1 indicates that all muscles within the hemisegment increased activity in the bend phase.

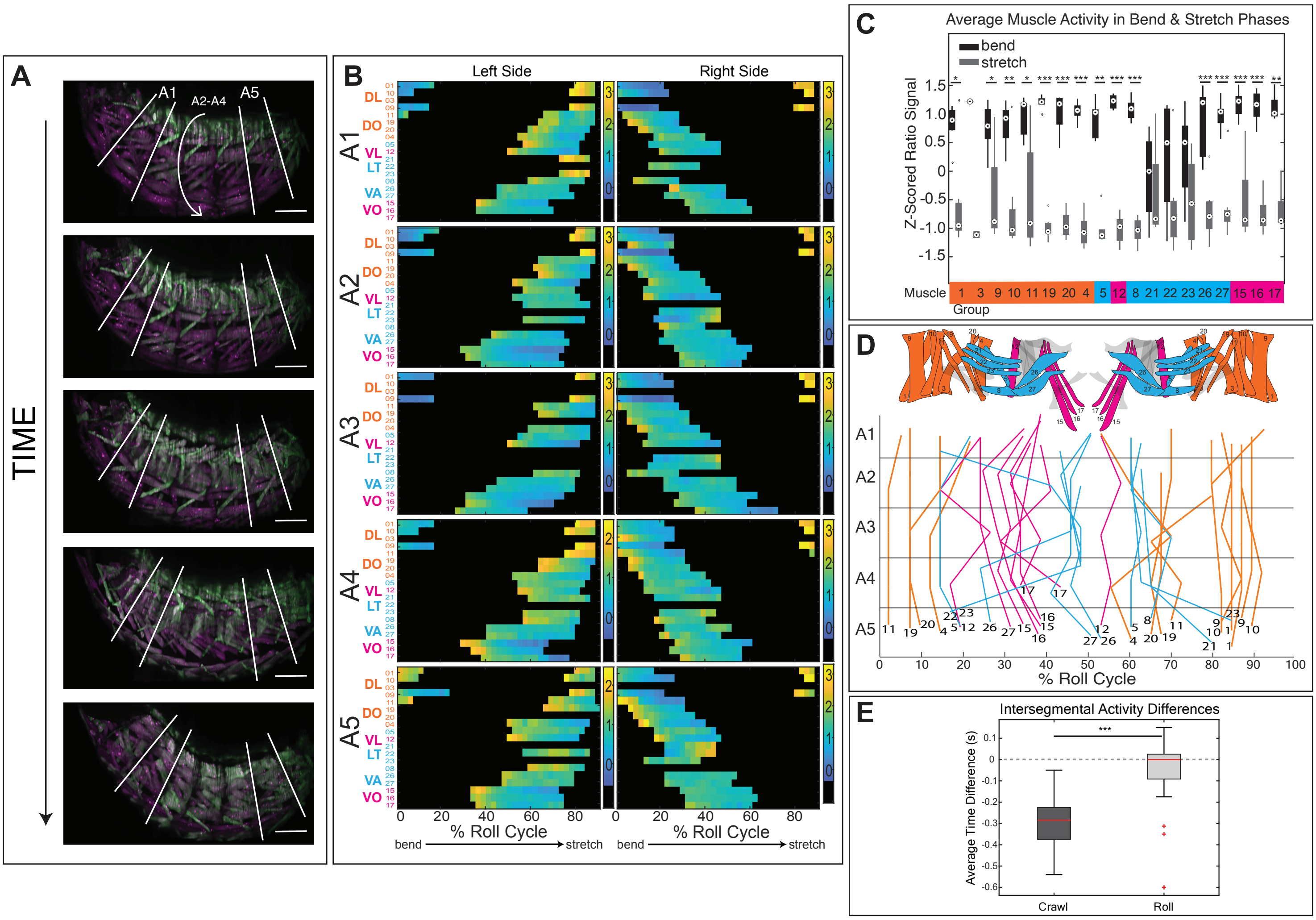
A
A1
A2-A4
A5
TIME
B
Left Side
Right Side
A1
A2
A3
A4
A5
DL
DO
VL
LT
VA
VO
% Roll Cycle
bend
stretch
C
Average Muscle Activity in Bend & Stretch Phases
bend
stretch
Z-Scored Ratio Signal
Muscle
Group
D
% Roll Cycle
E
Intersegmental Activity Differences
Average Time Difference (s)
Crawl
Roll

**Figure S3: Distal abdominal segments demonstrate similar muscle activity patterns to abdominal midsegments. (A**) SCAPE dual-color stills from single roll bout illustrating similar muscle GCaMP increases to midsegments in all segments. A1 and A5 are annotated and quantitatively analyzed in subsequent figure panels. **(B)** Heatmaps of z-scored ratiometric signal across all measured muscles in single roll bout, divided by segment (A2-A4) and side (left and right). Each row for each heatmap panel is an individual muscle, organized from dorsal (top row) to ventral (bottom row), and color-coded according to muscle groupings, as in **Figure 2, Figure S2**. Colorbars to the right of the upper heatmaps show range of ratiometric values. **(C)** Boxplot showing mean z-scored ratio signal for individual muscles from frames when muscles were along the bent side of the larva (black) vs. along the stretched side of the larva (gray) (*right*). Data are grouped and color-coded along x-axis as in **Figure 2** (n = 2 rolls, 2 larvae, 326 muscles). **(D)** Schematic of single segment with muscle color-coded by muscle grouping. Highest measured ratiometric muscle fluorescence times during single representative roll of extended segment analyses with individual muscle number labels. **(E)** Comparison between time difference of muscles in segments A2-A4 for forward crawling versus A1-A5 of rolling (crawl: n = 2 crawls, 2 larvae, 86 muscles; roll: n = 2 rolls, 2 larvae, 326 muscles). Negative values indicate that muscles in the adjacent posterior segment are active before muscles in the adjacent anterior segment and "0" indicates synchronous contraction. Mann-Whitney U tests were performed between intersegmental roll lag values and intersegmental crawl lag values. P values are indicated as ***$p<0.001$. Scale bars = 100µm (B).

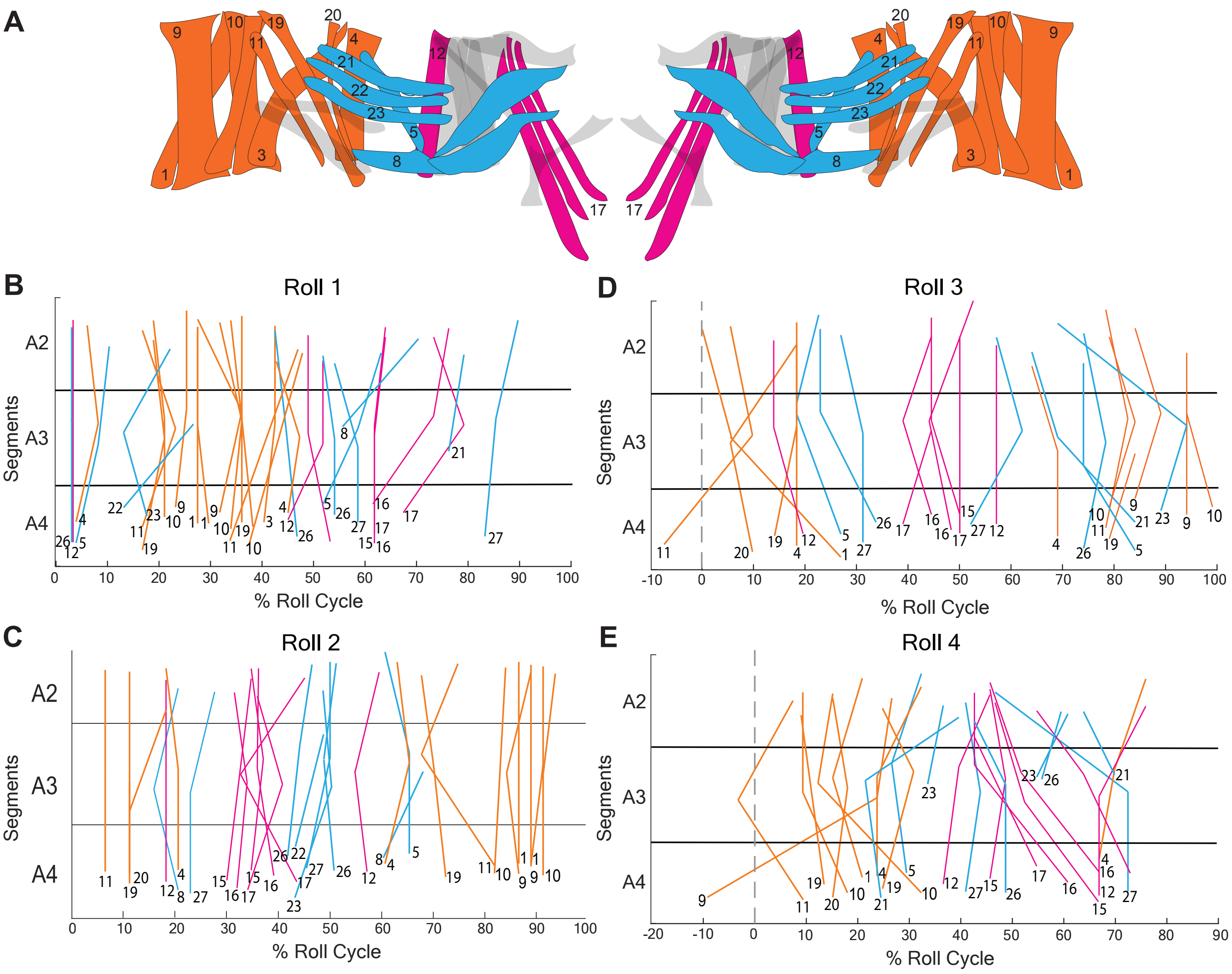


**Figure S4: Muscle activities occur as segmentally synchronous dorsoventral propagation. (A)** Segment color-coded by muscle group and with individual muscle numbers that correspond with color-coded muscle activity peak lines in subsequent panels. **(B-E)** Highest measured ratiometric muscle fluorescence times during single rolls (n = 4 rolls, 4 larvae). Note that, because rolling is a cyclical behavior, axes were adjusted in panels **D,E** to illustrate relative synchrony in segments where select muscles that appear twice during a single roll have offset peaks in linear time of a single roll.

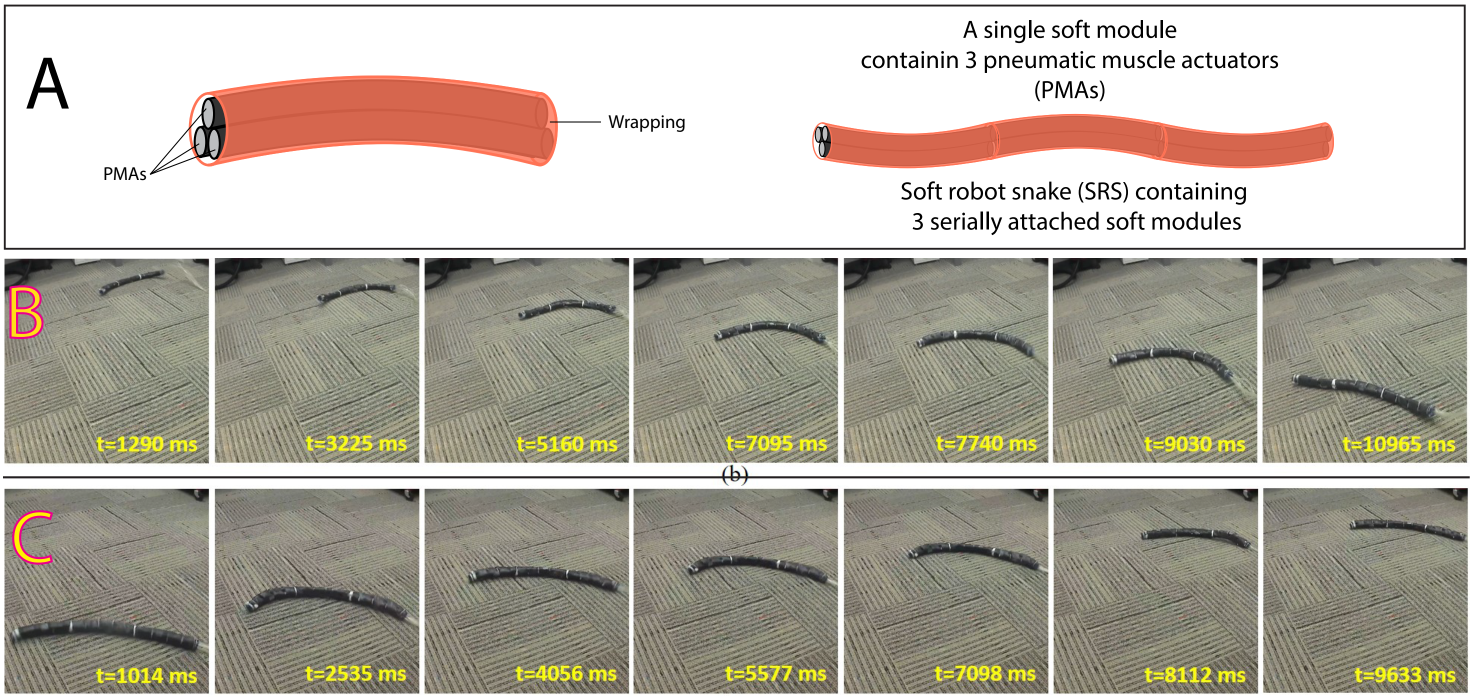


**Figure S5: Soft robot snake (SRS) inward and outward rolling locomotion identical to *Drosophila* larval rolling**. (Used with permission from Arachchige et al.)

A) The SRS developed by Arachchige et al. contains three serially attached segments of snake-like soft modules. Each segment contains 3 pneumatic muscle actuators (PMAs) triangularly fixated around the long axis of the snake (making it a total of 9 PMAs in the entire SRS), similar to larval longitudinal muscles. The length of each PMA is independently controlled by changing its air pressure through pneumatic supply lines, resembling muscle contraction and relaxation.
B) SRS inward rolling on a carpeted floor. The SRS is placed on the floor, bends and rolls toward the curve opening. C) SRS outward rolling. The robot bends and rolls away from the bend (opposite to the curve

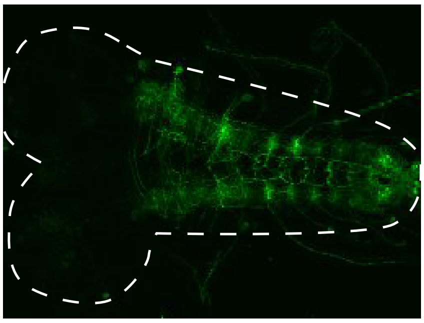

CQ>GtACR-EGFP

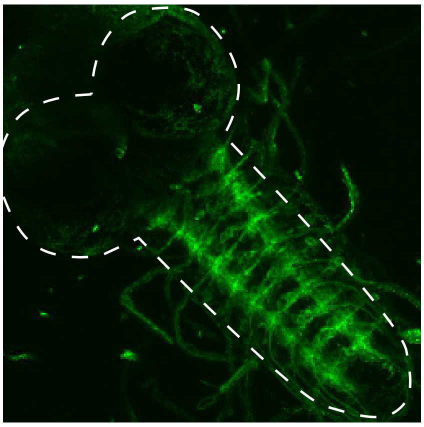

vGlut4AD-Nkx6DBD>
GtACR-EGFP

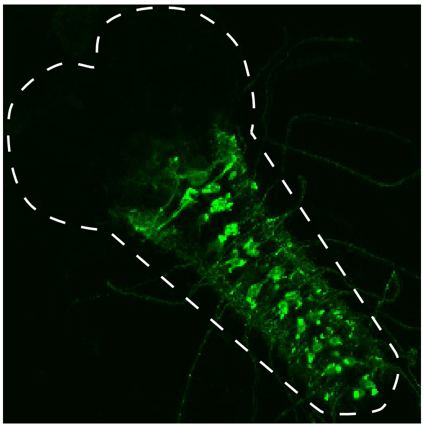

27E09>GtACR-EGFP

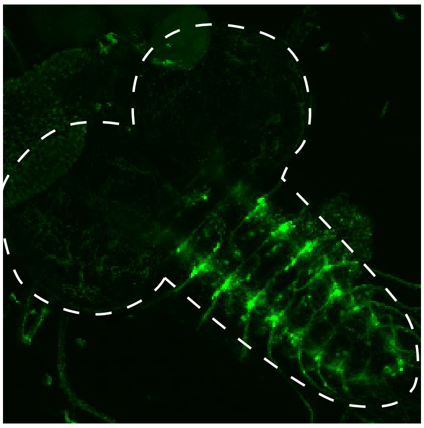

unc4AD-vGlutDBD>
GtACR-EGFP

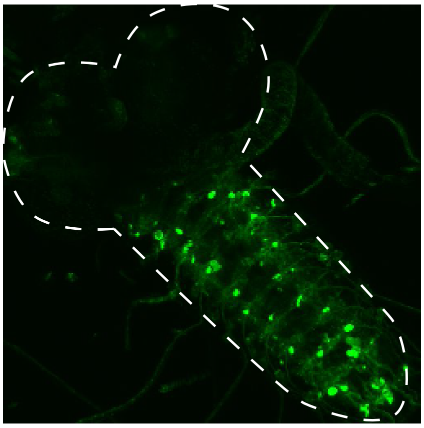

HB9>GtACR-EGFP

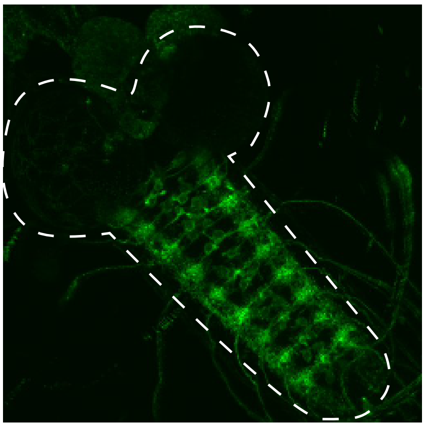

RRa>myr-EGFP

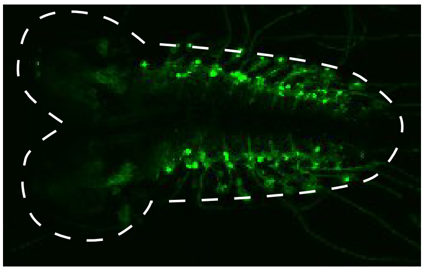

BH1>GtACR-EGFP

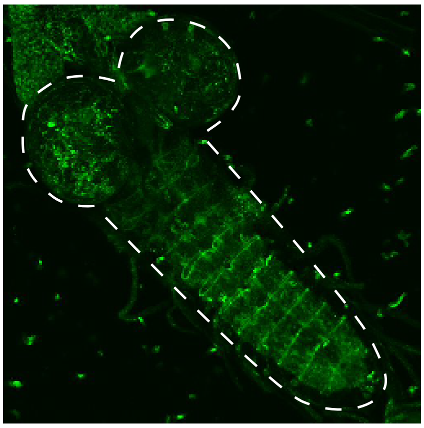

per>GtACR-EGFP

**Figure S6:** Expression pattern of UAS-GtACR1-eGFP in larval central nervous system (CNS) under the control of Motor Neuron(MN)- or per-Gal4 lines (*CQ>UAS-GtACR1-eGFP*, *Unc4$^{AD}$-vGlut$^{DBD}$>UAS-GtACR1-eGFP*, *HB9>UAS-GtACR1-eGFP*, *vGlut4$^{AD}$-Nkx6$^{DBD}$>UAS-GtACR1-eGFP, BH1> UAS-GtACR1-eGFP, 27E09> UAS-GtACR1-eGFP, per> UAS-GtACR1-eGFP; for RRa-Gal4, UAS-myr-GFP* is used for this imaging). White dashed lines show the outline of the isolated CNS, with two round brain lobes and a VNC extending outward. All *MN-Gal4>UAS-GtACR1-eGFP* shows few or no brain off-targets, and eGFP is seen in axons of MNs projecting out of the VNC, suggesting efficient GtACR1-eGFP expression. In *per>UAS-GtACR1-eGFP*, there are GtACR1-eGFP-positive glutamatergic interneurons in the VNC and in the brain. All images are max-intensity projections of larval CNS Z-stacks obtained from a Zeiss LSM-900 confocal microscope.

44H10::GCaMP6f (wild type) Forward crawling

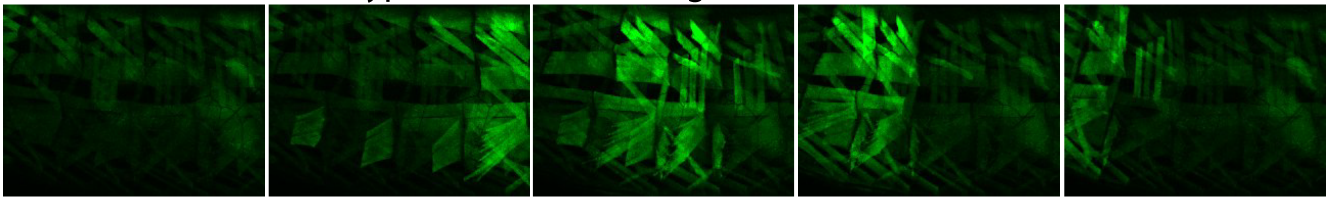

CQ>GtACR1 44H10::GCaMP6f Forward crawling

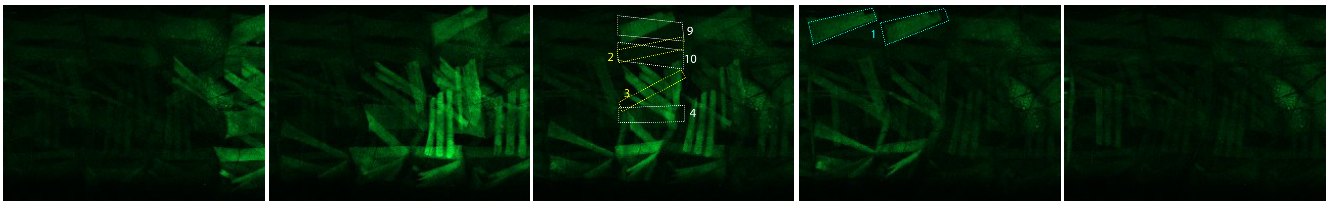

HB9>GtACR1 44H10::GCaMP6f Forward crawling

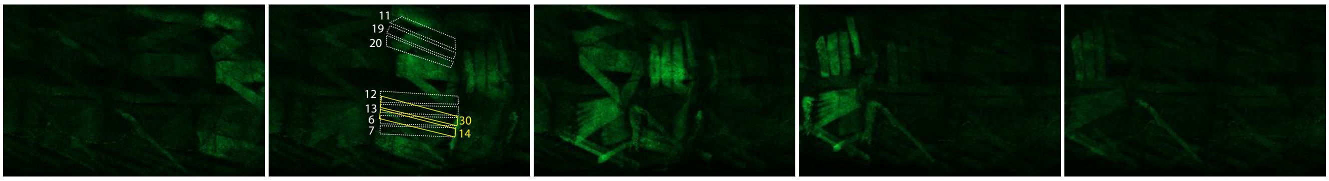

BH1>GtACR1 44H10::GCaMP6f Forward crawling

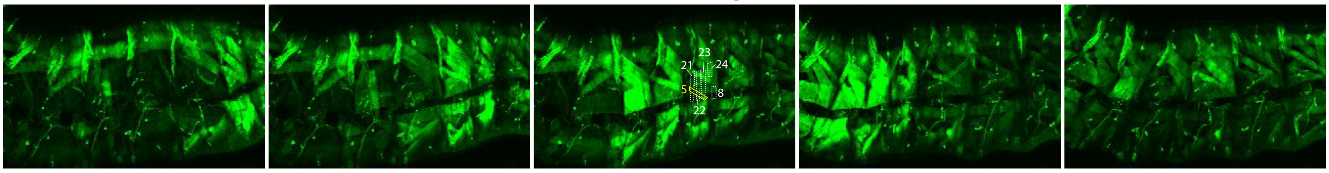

$unc4^{AD}$-$vGlut^{DBD}$ >GtACR1 44H10::GCaMP6f Forward crawling

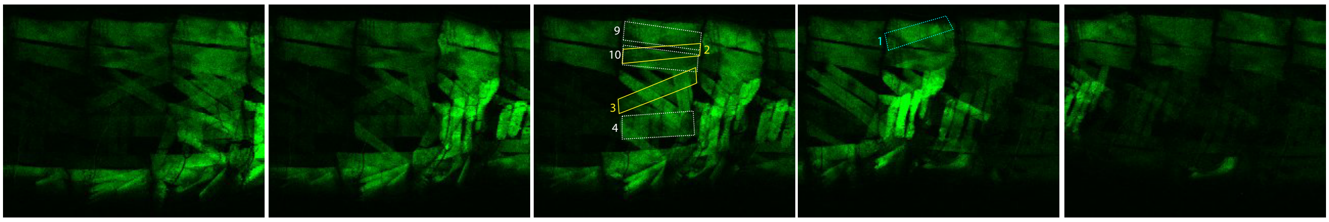

$vGlut4^{AD}$-$Nkx6^{DBD}$ >GtACR1 44H10::GCaMP6f Forward crawling

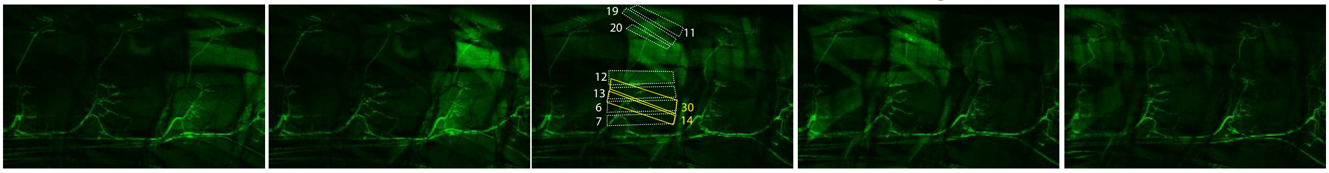

R27E09 + $unc4^{AD}$-$vGlut^{DBD}$ >GtACR1 44H10::GCaMP6f Forward crawling

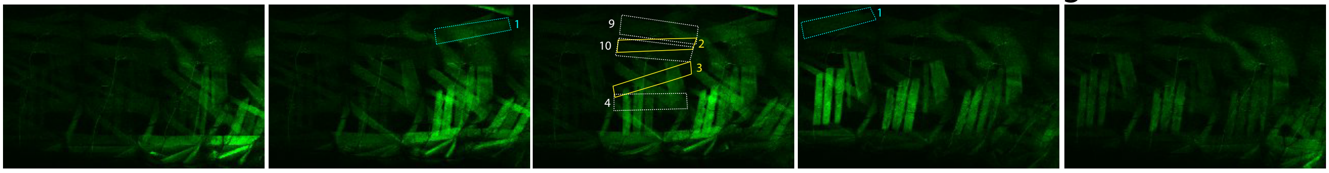

CQ+RRa>GtACR1 44H10::GCaMP6f Forward crawling

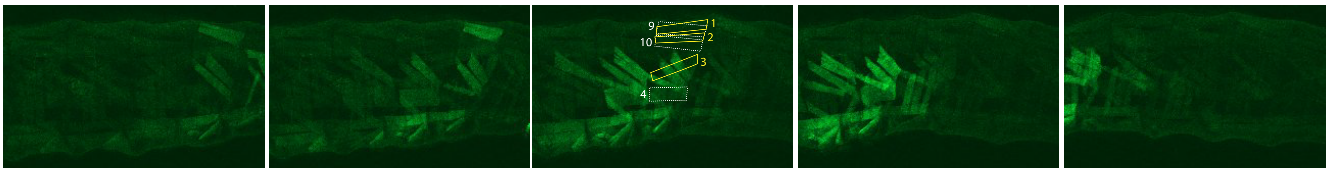

Time

**Figure S7:** Muscle GCaMP imaging during forward locomotion in intact larvae showing that GtACR1-eGFP sufficiently silences target muscles of motor neuron (MN) driven by MN-Gal4 lines. Sequential images of larvae with GCaMP6f calcium indicator in body wall muscles are shown. The genotype for control larvae is *44H10::GCaMP6f,UAS- GtACR1-eGFP*, and each *MN>UAS-GtACR1-eGFP* group also carries and the corresponding MN-Gal4 or MN-split-Gal4.

In control larvae, segments of body wall muscles are recruited sequentially, and muscle activity propagates toward the anterior (left). Every muscle in each segment becomes active once in each crawling bout.

In *CQ>UAS-GtACR1-eGFP* larvae (MN Ib silencing of muscles 2,3,4,9,10, shown in white or yellow boxes), the target muscles show little or no increase in GCaMP signal, suggesting *CQ>UAS-GtACR1-eGFP* completely silences muscle activity. In comparison, muscle 1 (cyan boxes) shows decent GCaMP activity.

*HB9>UAS-GtACR1-eGFP*, *vGlut4*$^{AD}$*-Nkx6*$^{DBD}$*>UAS-GtACR1-eGFP* (MN Ib silencing of muscles 11,19,20,12,13,6,7,30,14, white or yellow boxes), and *BH1>UAS-GtACR1-eGFP* (MN Ib silencing of muscles 21,22,23,24,5,8, white or yellow boxes) also show complete muscle silencing.

*unc4*$^{AD}$*-vGlut*$^{DBD}$*> UAS-GtACR1-eGFP* has partial muscle silencing of target muscles 2,3,4,9,10 (white or yellow boxes), meaning muscle activity is diminished but not fully abolished, compared to the non-target muscle 1 (cyan box). However, *unc4*$^{AD}$*-vGlut*$^{DBD}$ + *R27E09>UAS-GtACR1-eGFP* (targeting MN Is in addition to the Ib MNs 2,3,4,9,10) completely silenced the five *unc4*$^{AD}$*-vGlut*$^{DBD}$ targets. *CQ+RRa>UAS-GtACR1-eGFP* (targeting dorsal MN Is and MN Ib for muscles 1,2,3,4,9,10) completely silenced all six Ib targets of this combination.

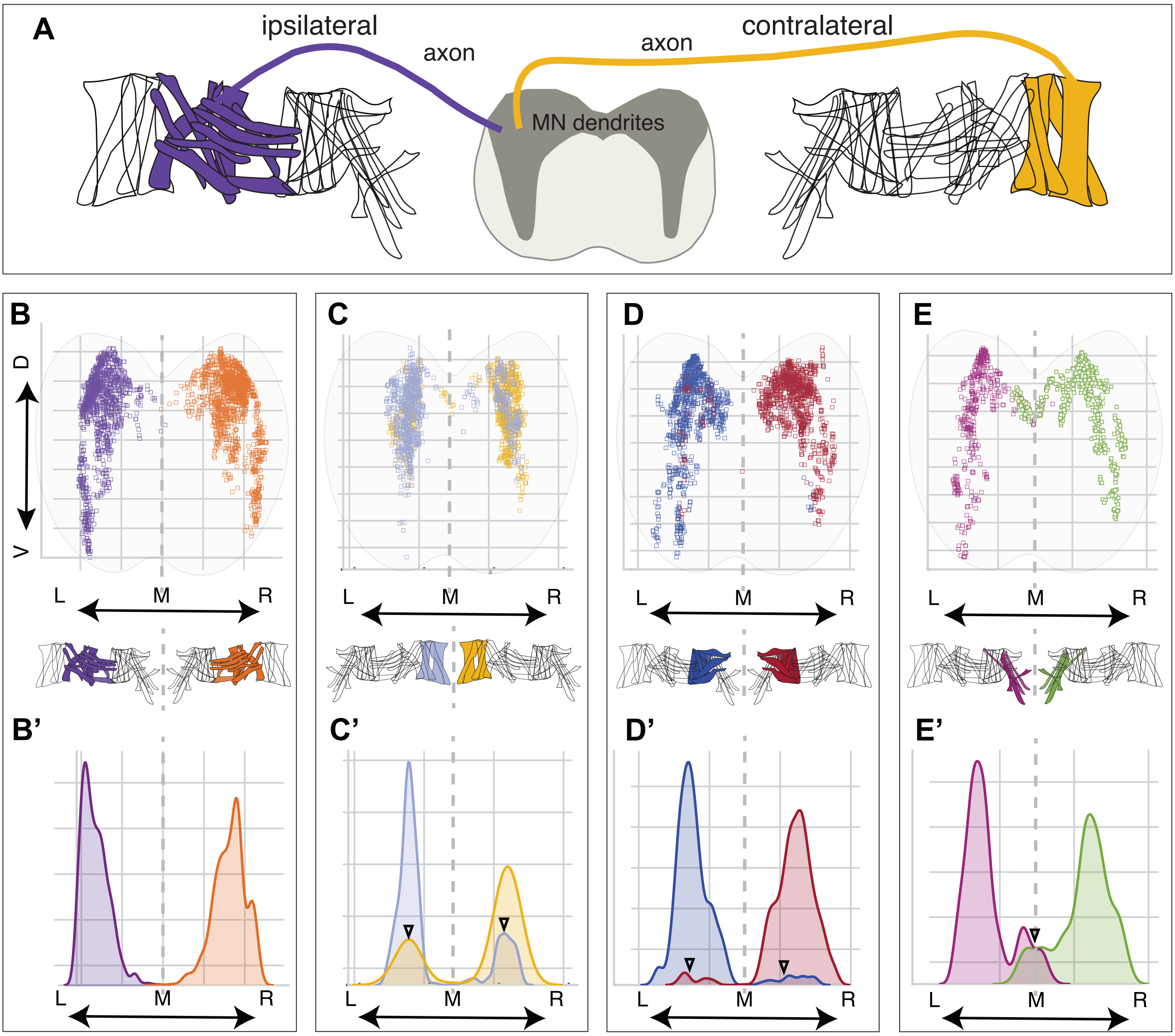
A
ipsilateral
axon
axon
contralateral
MN dendrites
B
D
V
L
M
R
B'
C
C'
D'
E
E'

**Figure S8: Left and right midline muscles receive inputs from partially overlapping regions of neuropil. (A**) Schematic showing motor neurons whose dendrites receive postsynaptic input either ipsilateral or contralateral to their axon projections. (**B-E**) Dorsoventral and mediolateral distributions of postsynaptic sites on MNs innervating the muscle groups indicated below each panel. Shaded gray area represents the cross-sectional view of the neuropil. Gray dashed lines define the midline (M) dividing the neuropil into left (L) and right (R) hemisegments, and the midline of dorsal and vental body wall muscles, respectively. (**B**) Postsynaptic sites of MNs innervating lateral muscles are localized exclusively in either left or right hemisegment of the neuropil. (**C-E**) Postsynaptic sites of MNs innervating muscles near the dorsal or ventral midlines occupy partially overlapping regions of the neuropil. (**B'-E'**) 1D kernel density plot of MN postsynaptic sites on the mediolateral axis. Black arrowheads in **C'-E'** show overlapping localization of postsynaptic sites of left and right MNs.

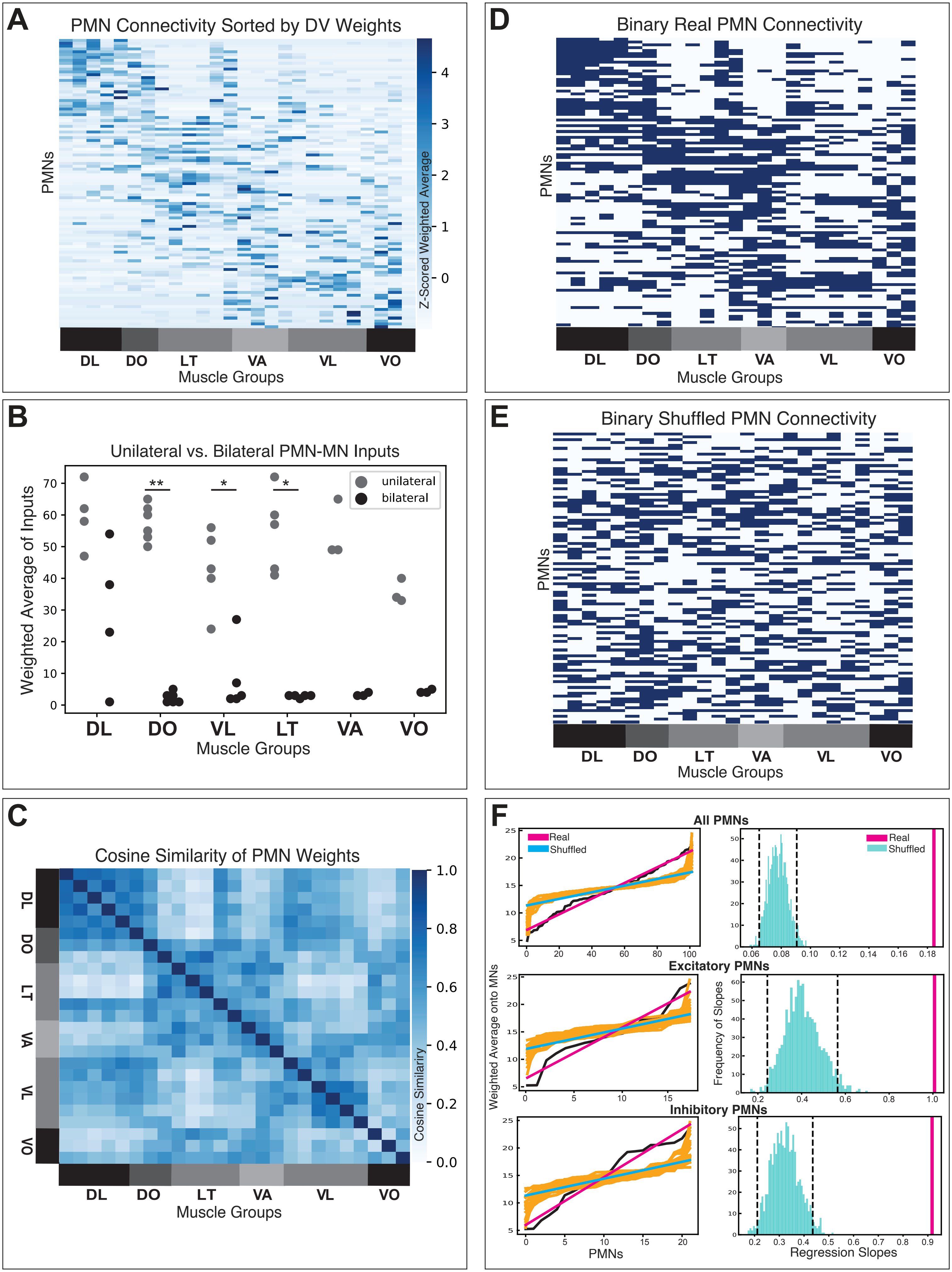

A
PMN Connectivity Sorted by DV Weights
PMNs
Z-Scored Weighted Average
DL DO LT VA VL VO
Muscle Groups
B
Unilateral vs. Bilateral PMN-MN Inputs
Weighted Average of Inputs
unilateral
bilateral
DL DO VL LT VA VO
Muscle Groups
C
Cosine Similarity of PMN Weights
Cosine Similarity
DL DO LT VA VL VO
Muscle Groups
D
Binary Real PMN Connectivity
PMNs
DL DO LT VA VL VO
Muscle Groups
E
Binary Shuffled PMN Connectivity
PMNs
DL DO LT VA VL VO
Muscle Groups
F
All PMNs
Real
Shuffled
Excitatory PMNs
Inhibitory PMNs
Weighted Average onto MNs
Frequency of Slopes
PMNs
Regression Slopes

**Figure S9: Premotor circuit organization supports circumferential muscle contraction sequence. (A**) Heatmap of synaptic weights of PMNs onto MNs, averaged across left and right PMNs and MNs and z-scored. PMNs (rows) are sorted by their weighted average onto MNs arranged dorsoventrally, illustrating that the PMN output onto MNs shows a global organization that permits muscle contraction in a dorsoventral sequence, as occurs in the circumferential muscle wave during rolling. (**B**). Quantitative comparison of PMN unilateral and bilateral projections onto MNs innervating specific spatial muscle subgroups. Each point represents the weighted average of PMN inputs onto a specific muscle within the muscle subgroup indicated on the x-axis. Mann-Whitney U tests were performed between weighted averages of PMNs onto unilaterally vs. bilaterally to MNs, demonstrating significant differences between unilateral vs. bilateral premotor innervation of muscle groups farther from the midline (DO, VL, and LT) and insignificant differences between muscle groups closer to the midline (DL, VA, and VO), providing a circuit structural correlate for premotor drive of circumferential muscle contractions in rolling. P values are indicated as $^*p<0.05$, $^{**}p<0.01$. (**C**) Cosine similarity of PMN inputs to MNs organized in dorsoventral order demonstrates that MNs innervating muscle groups that are closer together in dorsoventral axis have a high degree of shared input from PMNs. (**D**) Binarized synaptic weights of PMNs onto MNs organized dorsoventrally, as in (**A**). (**E**) Example of shuffled binary synaptic weights from PMNs onto MNs, where connections from PMNs onto specific MNs were randomly selected while preserving the total number of outputs from each PMN and the probability of input onto each MN. Notably, the shuffling procedure, despite preserving global statistics of PMN outputs and MN inputs, noticeably removes the dorsoventral preferences of real PMNs observed in (**A**) and (**D**). (**F**) (Left) Quantitative comparison of weighted averages of PMNs onto MNs arranged in dorsoventral order from binary real PMN-MN connectome data (as in (**D**)) vs. 1000 binary shuffled PMN-MN connectivity matrices (as in example (**E**)). Real weighted average of PMNs onto MNs for all PMNs (top), PMNs previously identified as excitatory (middle), and PMNs previously identified as inhibitory (bottom) is shown in black, with corresponding regression line in magenta. Weighted averages from 1000 randomly shuffled PMN-MN matrices shown in gold, with corresponding mean regression line across all shuffles in cyan. Regression lines of real PMN-MN weighted averages for all PMNs, excitatory-only PMNs, and inhibitory-only PMNs demonstrate steeper slopes (higher regression coefficients) than that of the shuffled PMN-MN weighted averages, highlighting their stronger dorsoventral spatial preference than given by chance. (Right) Histograms illustrating frequency of regression coefficients over 1000 shuffled PMN-MN matrices (cyan). Real-valued regression coefficient (magenta) falls well outside of the 95% confidence bounds of the distribution of shuffled regression coefficients (black dashed lines), indicating that dorsoventral structure of PMN-MN connectivity is significantly different from chance.

A02h

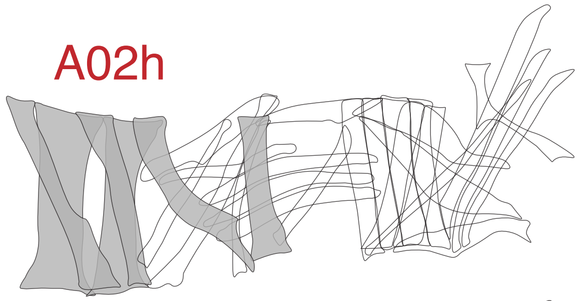

A02m/n_a1r

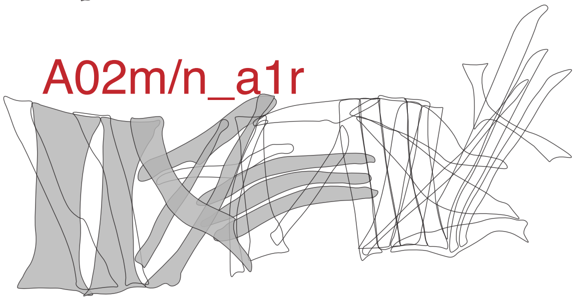

A02f

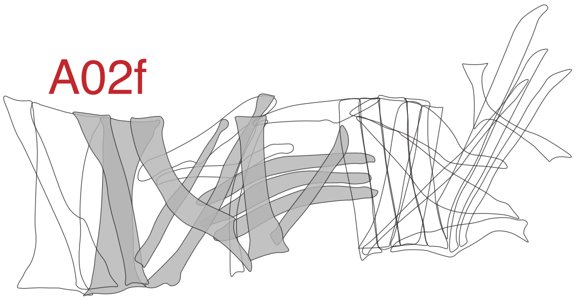

A02e

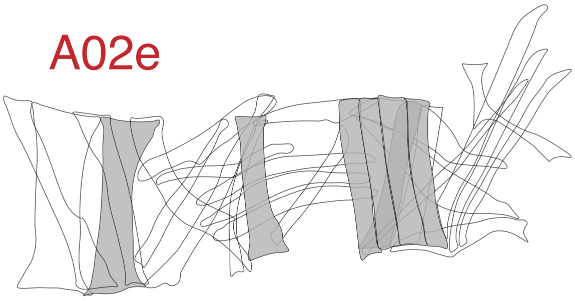

A02i

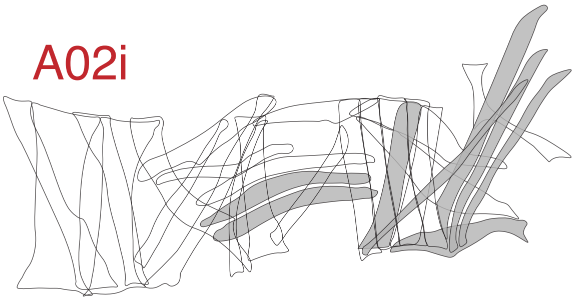

A02g

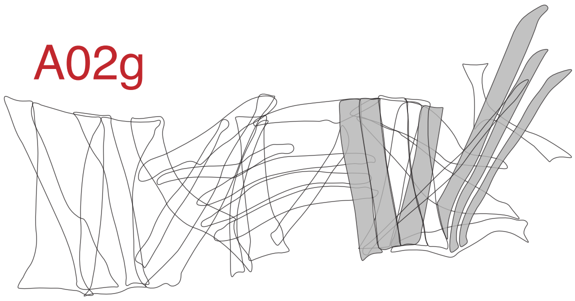

**Figure S10:** A02 PMNs are glutamatergic PMNs that synapse onto MNs. Here the target muscles of MNs receiving inputs from A02 PMNs are shown in gray. Each PMN synapses with MNs innervating neighboring groups of muscles in dorsal, dorsolateral, ventrolateral, or ventral areas of body wall muscles. We propose these PMNs contribute to circumferential progression of muscular activity during rolling.
